# Mitochondria-ER contacts restrain store-operated Ca²⁺ entry via Ca²⁺ flickering and STIM1 trapping

**DOI:** 10.1101/2025.01.17.633482

**Authors:** Yu-Chiao Lin, Chieh-Ju Sung, Yu-Chun Lin, Hsuan-Chao Lin, Pei-Ju Tsai, Mei-Leng Cheong, Feng-Chiao Tsai

## Abstract

Ca²⁺ homeostasis requires coordinated regulation of ER stores and mitochondrial buffering. While the ER replenishes Ca²⁺ via STIM1-mediated store-operated Ca²⁺ entry (SOCE), how mitochondria–ER contact sites (MERCs) influence this process remains unclear. Here, using fluorescent live imaging, we capture spontaneous mitochondrial Ca²⁺ flickering at MERCs, which is driven by the IP₃R–VDAC1-mediated Ca²⁺ transfer and exerts an inhibitory effect on SOCE. Mechanistically, MERCs establish local ER Ca²⁺ depletion microdomains through this Ca²⁺ transfer. These microdomains act as molecular traps, triggering STIM1 accumulation at MERCs via its polybasic K-domain and sequestering it away from the plasma membrane (PM) to suppress SOCE. Furthermore, acute MERC induction redistributes constitutively active, Ca²⁺-insensitive STIM1 away from ER–PM junctions, demonstrating that MERCs directly outcompete the PM for STIM1 recruitment. Finally, the microtubule (EB1)–ER contacts serve as a finite-capacity reservoir that buffers excess STIM1, with its disruption unmasking the dynamic competition between MERCs and PM. Together, our findings establish a tripartite system—PM, mitochondria, and microtubules—that dictates SOCE by controlling STIM1 topography, thereby protecting cells against Ca²⁺ overload.

## Main text

Ca^2+^ signaling is essential for proper biological functions ranging from gene expression to cell death (Clapham, 2007; La Rovere *et al*, 2016) through the spatial and temporal control of Ca^2+^ flows. As the primary intracellular Ca^2+^ reservoir, the endoplasmic reticulum (ER) Ca^2+^ levels are dynamically regulated by the coordination of sarcoplasmic/endoplasmic reticulum Ca^2+^ ATPases (SERCAs) and inositol 1,4,5-trisphosphate receptors (IP_3_Rs). Upon ER Ca^2+^ depletion, cells activate store-operated Ca^2+^ entry (SOCE) – a process essential not only for replenishing ER stores but also for sustaining the cytosolic Ca^2+^ elevations required for a broad spectrum of physiological responses, including gene expression, secretion, and cell proliferation. In response to store depletion, the ER-resident sensor stromal interaction molecule 1 (STIM1) oligomerizes, translocates to ER-plasma membrane (PM) junctions, and activates ORAI1 channels for Ca^2+^ entry (Liou *et al*, 2005; Park *et al*, 2009; Prakriya *et al*, 2006; Roos *et al*, 2005; Vig *et al*, 2006; Wu *et al*, 2006). While ER Ca^2+^ depletion serves as the master trigger for this process, STIM1 activation is also fine-tuned by regulatory proteins, post-translational modifications, and cytoskeletal elements including microtubules (Chang *et al*, 2018; Jing *et al*, 2015; Palty *et al*, 2012).

Functioning as high-capacity Ca²⁺ buffers for intracellular signaling, mitochondria regulate Ca²⁺ influx through the inner-membrane mitochondrial Ca^2+^ uniporter (MCU) (Baughman *et al*, 2011; De Stefani *et al*, 2011; Kirichok *et al*, 2004) and the outer-membrane voltage-dependent anion channels (VDACs)(Colombini, 2004), while Ca²⁺ efflux is mediated by the mitochondrial Na^+^/Ca^2+^ exchanger (NCLX) (Palty *et al*, 2010) and the Leucine Zipper-EF-Hand Containing Transmembrane Protein 1 (LETM1) (Austin *et al*, 2017; Jiang *et al*, 2009). Because resting cytosolic Ca^2+^ levels are insufficient to drive MCU-mediated Ca^2+^ uptake, efficient mitochondrial Ca^2+^ influx depends on the close physical apposition (10–30 nm) maintained at mitochondria–ER contact sites (MERCs)(Csordas *et al*, 2006; Csordas *et al*, 2010). The structural integrity of MERCs is sustained by tethering proteins, including mitofusin 2 (MFN2)(de Brito & Scorrano, 2008; Naon *et al*, 2023), PDZD8 (Hirabayashi *et al*, 2017), FUNDC1 (Wu *et al*, 2017; Wu *et al*, 2016) and the VAPB-PTPIP51 complex (De Vos *et al*, 2012). These tethers coordinate with the IP₃R–GRP75–VDAC1 channeling complex (Szabadkai *et al*, 2006) to facilitate the localized Ca^2+^ transfer required for mitochondrial bioenergetics and cell survival. Early work established that this spatially confined transfer generates localized mitochondrial Ca²⁺ signals, including miniature transients driven by ryanodine receptors (Pacher *et al*, 2002) and ER-to-mitochondria Ca^2+^ transfer events at MERCs (Cardenas *et al*, 2020; Cardenas *et al*, 2010; Csordas *et al*., 2010; Duchen *et al*, 1998; Hou *et al*, 2013; Rizzuto *et al*, 1998). However, whether and how these local Ca²⁺ events reciprocally regulate global Ca²⁺ homeostasis remains unclear.

Although STIM1 is classically defined by its function at ER–PM junctions, emerging evidence highlights its potential role at MERCs. Proximity-labeling proteomics (Cho *et al*, 2020) and interactome profiling (Sanchez-Lopez *et al*, 2024) have identified STIM1 as a key interactor of MERC-resident proteins, including GRP75 and PTPIP51. This work recently culminated in the landmark discovery that a dedicated pool of STIM1 localizes to MERCs, where it facilitates direct ER-to-mitochondria Ca^2+^ flux through interaction with GRP75 (Orantos-Aguilera *et al*, 2026). These findings raise a fundamental question: how do localized mitochondrial Ca²⁺ dynamics at MERCs—and the STIM1 subpopulation residing there—influence the canonical translocation of STIM1 to the plasma membrane? Whether these distinct spatial roles of STIM1 represent independent functions or a coordinated regulatory trade-off remains a central, unresolved question.

Here, using fluorescent live imaging, we identify MERCs as pivotal spatial regulators of SOCE that orchestrate Ca²⁺ homeostasis through competitive STIM1 sequestration. We show that IP₃R–VDAC1-mediated Ca²⁺ transfer generates mitochondrial Ca²⁺ flickers while establishing local ER Ca²⁺ depletion microdomains. These microdomains trap STIM1 at MERCs via its polybasic domain, thereby suppressing its translocation to the plasma membrane. We further demonstrate that MERC induction actively redistributes pre-formed STIM1 clusters away from ER-PM junctions, revealing a dynamic and competitive remodeling of STIM1 topography.

Finally, we establish that the microtubule (EB1)–ER contacts function as a finite-capacity STIM1 reservoir that competes with MERCs and ER-PM junctions for the available STIM1 pool. Collectively, our findings uncover an integrated tripartite regulatory system comprising the PM, mitochondria, and microtubules. This system fine-tunes SOCE via competitive STIM1 sequestration to protect cells against Ca²⁺ overload.

## RESULTS

### MCU-mediated mitochondrial Ca^2+^ flickers occur spontaneously at MERCs

To investigate basal mitochondrial Ca^2+^ dynamics, we performed live-cell imaging using the sensitive, intensiometric Ca^2+^ indicator mito-LAR-GECO1.2 in HeLa and HepG2 cells. Under resting conditions, individual mitochondria exhibited spontaneous, transient Ca^2+^ elevations restricted to single mitochondria or small clusters within the network—events we term “mitochondrial Ca^2+^ flickering” (**Figure 1A, 1B** and **Video S1**). To exclude artifactual signals from mitochondrial motility, we co-expressed the matrix-targeted marker mito-GFP as a movement reference. The Ca^2+^ flickering events persisted independently of positional changes (**Figure 1C** and **Video S2**). Sensor responsiveness was confirmed by SERCA inhibitor thapsigargin (TG) and ionomycin (**Figure S1A**). Quantitative analysis in HepG2 cells revealed that these flickers occur at a frequency of ∼ 1.5 ± 0.5 min^-1^ per mitochondrion with amplitudes ranging from 0.9 to 1.3-fold of average signals (mean ± s.d. = 1.1 ± 0.1 fold) (**Figure 1D**). Given that basal mitochondrial Ca^2+^ levels typically reside between 100-200 nM (David *et al*, 2003; Rizzuto *et al*, 1992), these Ca^2+^ flickers are estimated to reach 90-260 nM in amplitudes. In addition, individual Ca^2+^ flickers lasted 6 to 33 seconds (mean ± s.d. = 19.4 ± 6.9 sec) (**Figure 1D**). Peak-aligned averaging demonstrated a rapid rise and decay phase, while autocorrelation analysis confirmed their non-periodic nature (**Figure 1E**). Critically, the mean flicker frequency equaled its variance, following a Poisson distribution (**Figure 1F**) and establishing each mitochondrial flicker as an autonomous stochastic unit operating with a fixed probability.

**Figure 1.**
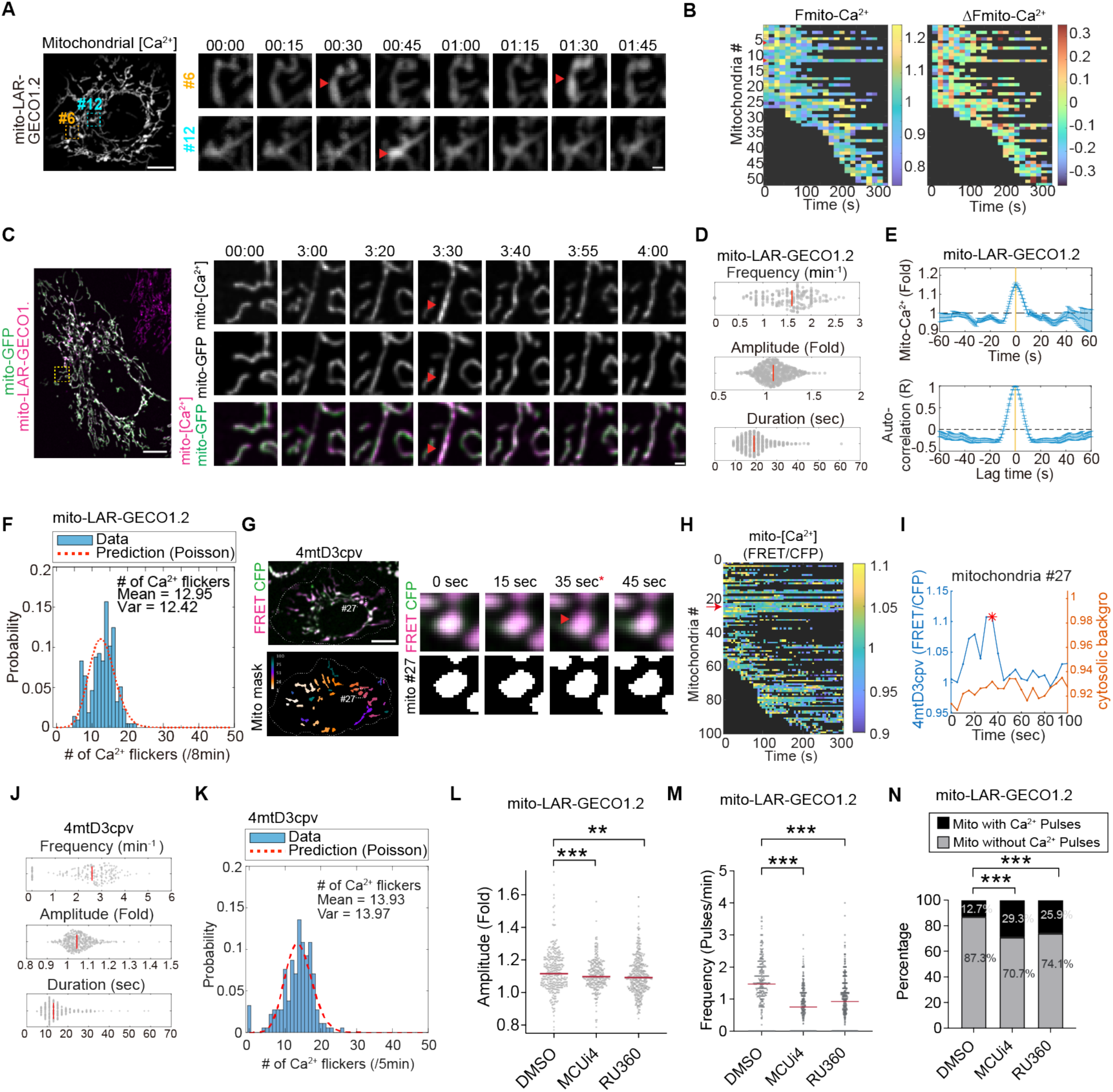
Mitochondrial Ca^2+^ flickers spontaneously and stochastically via MCU-dependent matrix uptake. **A.** Representative time-lapse images of HeLa cells expressing the mitochondrial Ca^2+^ sensor mito-LAR-GECO1.2. Red arrowheads indicate spontaneous, localized transient increases in Ca^2+^ signal (flickers) in individual mitochondria (e.g., #6 and #12 corresponding to the indices in panel **B**). **B.** Heatmaps illustrating the spatiotemporal dynamics of mitochondrial Ca^2+^ flickers. The left panel shows the normalized fluorescence (F/F_mean_) of individual mitochondria (indexed on the y-axis) plotted over time. F_mean_ represents the time-averaged fluorescence intensity for each respective mitochondrion. The right panel displays the corresponding frame-to-frame fluorescence change (ΔF = F_t_ − F_t−1_) to highlight the occurrence and amplitude of the transient events. **C.** Dual-color imaging of mitochondria expressing both mito-GFP (green) and a red mitochondrial Ca^2+^ sensor (magenta). The time-lapse sequence confirms that the transient fluorescence spikes represent biological Ca^2+^ fluctuations rather than mitochondrial movement or morphological artifacts. **D.** Distribution analysis of mitochondrial Ca^2+^ flicker properties showing frequency (events per mitochondrion; top), peak amplitude (fold change relative to baseline of each flicker event, middle), and duration (seconds; bottom). Red vertical lines indicate median values. **E.** Temporal characterization of Ca²⁺ flickers. Top: Peak-aligned average trace of Ca^2+^ transients, demonstrating their non-oscillatory nature. Bottom: autocorrelation analysis, confirming a lack of periodicity of these Ca^2+^ events. **F.** Frequency distribution of mitochondrial Ca^2+^ flickers fitted to a Poisson distribution. The histogram displays the probability of specific flicker numbers per mitochondrion over an 8-minute recording period. The red dashed line represents the Poisson fit (Mean = 12.95, Variance = 12.42), indicating that the flickers are stochastic and independent events. **G.** Validation of spontaneous flickers using a FRET-based mitochondrial Ca^2+^ sensor, 4mtD3cpv. Top left: representative pseudo-colored FRET/CFP images. Bottom left: masks with colors indexing individual mitochondria. Right: Time-lapse FRET/CFP signals of mitochondrion #27, highlighting a flicker event (red arrowhead) at 35 sec. **H.** Spatiotemporal heatmap of FRET/CFP signals corresponding to panel G. Red arrow indicates mitochondrion #27. **I.** Representative FRET/CFP trace of mitochondrion #27 (blue) compared to the cytosolic background signal (orange), demonstrating that the flicker (red asterisk) is a discrete mitochondrial event. **J.** Beeswarm plots showing the distribution of flicker frequency, amplitude, and duration measured by the 4mtD3cpv sensor. Red lines indicate the median. **K.** Probability histogram of flicker occurrence per mitochondrion over a 5-min recording period using 4mtD3cpv. Flickers are fitted with a Poisson distribution (red dashed line; Mean = 13.93, Variance = 13.97), indicating their stochastic nature. **L.** Amplitude of mitochondrial Ca^2+^ flickers in cells treated with DMSO (control) or the MCU inhibitors MCUi4 and RU360. Beeswarm plots show the distribution of flickers amplitude with red line indicates the median. **M.** Frequency of mitochondrial Ca^2+^ flickers in the corresponding treatment groups. Beeswarm plots show the distribution of flickers amplitude with red line indicate the median. **N.** Bar chart quantifying the percentage of mitochondria exhibiting flicker activity under the indicated treatments. Statistical significance was determined by an unpaired, two-tailed Student’s t-test for all panels except for Figure 1N, which was analyzed using a Chi-square test. (*P < 0.05; **P < 0.01; ***P < 0.001). Scale bars: 10 μm; inset, 1 μm.

To rule out probe-specific artifacts, we independently validated those observations using the FRET-based ratiometric sensor 4mtD3cpv. Heatmaps and single-organelle traces confirmed the presence of asynchronous flickers (**Figure 1G–I**), with kinetic signatures and Poisson statistics indistinguishable from those obtained with mito-LAR-GECO1.2 (**Figure 1J**, **1K** and **S1B**), collectively substantiating the physiological reality of mitochondrial Ca^2+^ flickers. To determine if these Ca^2+^ events were driven by transient oxidative bursts—previously recorded as spontaneous “redox flashes” (Wang *et al*, 2008) and proposed co-regulators of Ca^2+^ signaling (Booth *et al*, 2021; Patel *et al*, 2022)—we utilized the ratiometric redox sensor matrix-roGFP, which revealed no oxidative fluctuations coinciding with Ca^2+^ flickers (**Figure S1C** and **S1D**). Furthermore, neither elevating mitochondrial ROS with the targeted redox cycler mitoParaquat (mitoPQ) (Robb *et al*, 2015) nor pharmacological ROS manipulation altered the spatiotemporal occurrence or kinetic properties of the flickers (**Figure S1E–G**), demonstrating that these events are ROS-independent.

Turning to the molecular machinery of Ca²⁺ influx, we focused on the mitochondrial calcium uniporter (MCU), which is the primary channel for Ca^2+^ uptake into the matrix. Since MCU has a low affinity for Ca^2+^, it remains mostly inactive during resting bulk cytosolic Ca^2+^ levels. Therefore, we investigated its role in mitochondrial Ca^2+^ flickering. Pharmacological inhibition of MCU using MCUi4 and RU360 significantly attenuated flicker amplitude (**Figure 1L**), frequency (**Figure 1M**), and the proportion of active mitochondria (**Figure 1N**), establishing MCU as the essential uptake mechanism into the matrix. Importantly, while these MCU-dependent flickering events serve as a direct optical reporter of active ER-to-mitochondria Ca²⁺ flux, this matrix accumulation is downstream of the local transport events occurring within the intermembrane space.

Thus, the flickering activity functions as a high-fidelity proxy for real-time contact site clearance, confirming the involvement of high-Ca²⁺ microdomains typically generated at MERCs.

To test whether MERCs are the source of this Ca²⁺ signal, we structurally uncoupled ER and mitochondria by knocking down the tethering proteins MFN2 (de Brito & Scorrano, 2008; Naon *et al*., 2023) and PDZD8 (Hirabayashi *et al*., 2017) via short hairpin RNAs (shMFN2 and shPDZD8). Knockdown efficiency was confirmed (**Figure S2A** and **S2B**), and 3D reconstructions verified a significant reduction in MERCs (**Figure 2A** and **2B**). Strikingly, MERC disruption profoundly suppressed spontaneous Ca²⁺ flickering: spatiotemporal heatmaps showed significant reduction of flicker activity (**Figure 2C**), with significant reductions in the proportion of active mitochondria (**Figure 2D**), flicker amplitude (**Figure 2E**) and frequency (**Figure 2F**), while event duration remained unchanged (**Figure S2C**). These findings were fully replicated using the 4mtD3cpv sensor (**Figure 2G–I** and **S2D**). Conversely, we directly induced MERCs formation using a rapamycin-inducible dimerization system, in which an ER-targeted FRB domain and an outer mitochondrial membrane-targeted FKBP are brought into proximity upon rapamycin addition, forcing ER-mitochondria apposition within 60 seconds (**Figure S2E**, **S2F** and **Video S3**). While physical ER-mitochondria tethering has been shown to facilitate steady-state Ca^2+^ transfer (Csordás et al., 2010), our results demonstrate that these contacts are sufficient to drive spontaneous, stochastic Ca^2+^ flickering. Using an acute rapamycin-inducible system, we observed that the formation of new MERCs immediately triggers localized Ca^2+^ transients (**Figure 2J, 2K**), establishing MERCs as the primary structural requirement for mitochondrial flickering dynamics. Together, these loss- and gain-of-function results confirmed that physical ER-mitochondria coupling at MERCs is both necessary and sufficient to drive spontaneous mitochondrial Ca²⁺ flickering.

**Figure 2.**
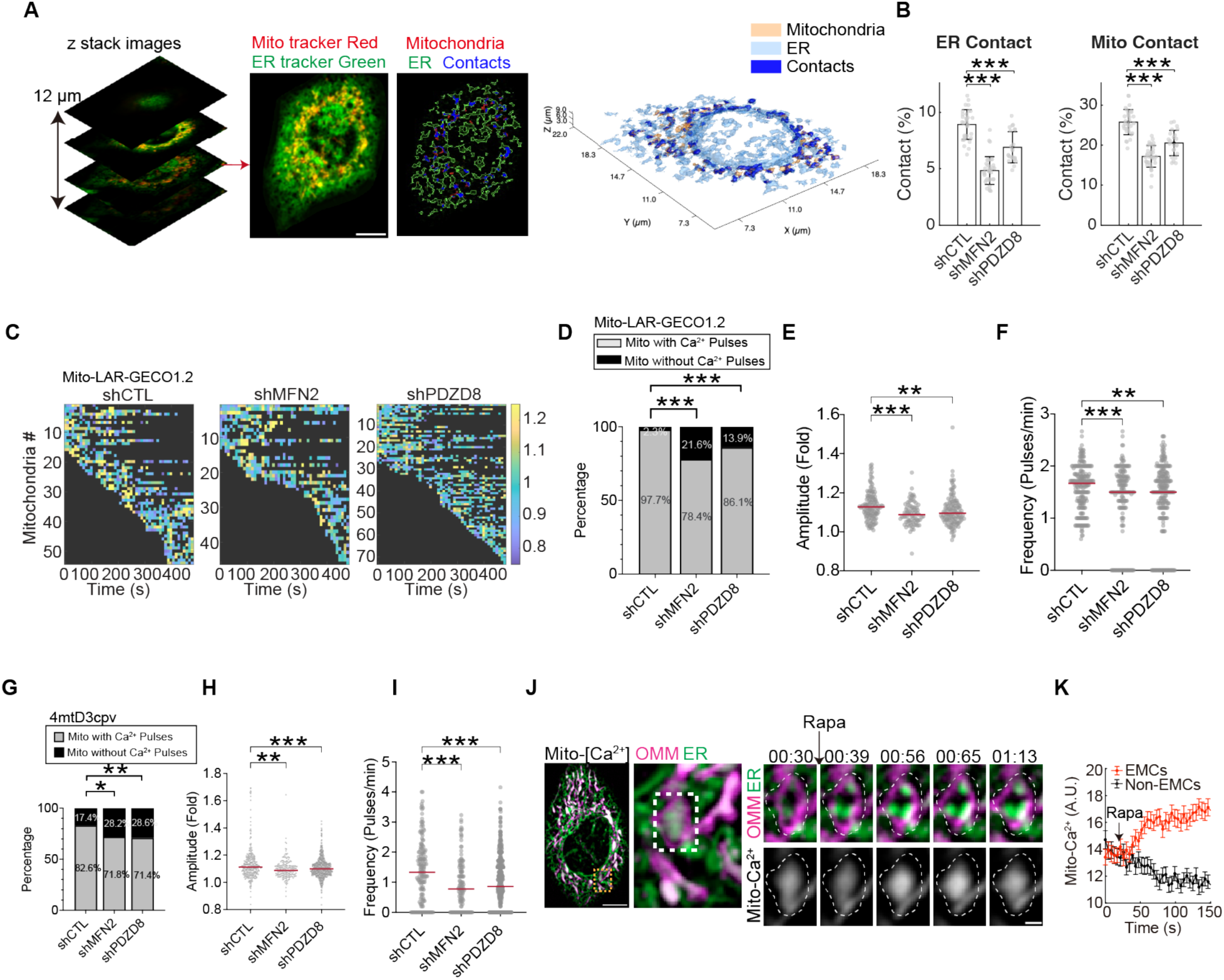
Mitochondrial Ca^2+^ flickers are driven by mitochondria-ER contact sites (MERCs). **A.** Representative confocal Z-stack imaging strategy for evaluating mitochondria-ER contact sites (MERCs). Images show HeLa cells co-stained with MitoTracker Red and ER-Tracker Green. The middle panel displays the 2D segmentation of mitochondria (red outlines), the endoplasmic reticulum (ER, green outlines), and their colocalized contact regions (blue). The rightmost panel represents a 3D surface rendering reconstructed from the Z-stack images. Scale bar, 10 μm. **B.** Quantification of the percentage of the ER surface in contact with mitochondria (ER contact, left) and the mitochondrial surface in contact with the ER (Mito contact, right) in control (shCTL) cells and cells following knockdown of the mitochondria-ER tethering proteins MFN2 (shMFN2) or PDZD8 (shPDZD8). **C.** Heatmaps illustrating the spatiotemporal dynamics of mitochondrial Ca^2+^ flickers in shCTL, shMFN2, and shPDZD8 HepG2 cells expressing the mito-LAR-GECO1.2 sensor. **D.** Bar chart quantifying the percentage of mitochondria exhibiting spontaneous Ca^2+^ flicker activity across the indicated groups. **E–F.** Quantitative analysis of mitochondrial Ca^2+^ flicker amplitude (E), frequency (F) in shCTL, shMFN2, and shPDZD8 cells. Red lines indicate the median. **G–I.** Validation of flicker properties using the FRET-based sensor 4mtD3cpv. Panels show the percentage of active mitochondria (**G**), and the distribution of flicker amplitude (**H**), frequency (**I**) in control and knockdown cells. Red lines indicate the median. **J.** Acute induction of synthetic MERCs drives localized mitochondrial Ca^2+^ uptake. Representative images show a mitochondrion (OMM, magenta) and the recruited ER (green) post-rapamycin treatment, with corresponding localized Ca^2+^ dynamics visualized via mito-LAR-GECO1.2 (grayscale). Dotted lines in the time-lapse insets demarcate a mitochondrion where MERCs were induced by rapamycin. The sequence highlights the emergence of discrete, flickering mitochondrial Ca^2+^ signals specifically at the mitochondrion with induced contact sites. Scale bar, 1 µm. **K.** Quantification of mitochondrial Ca^2+^ fluorescence kinetics at newly formed contact regions (MERCs, red trace) compared to adjacent, non-tethered regions (Non-MERCs, grey trace), corresponding to the condition shown in (J). Statistical significance was determined by an unpaired, two-tailed Student’s t-test assuming equal variance for all panels except for Figure 2D and 2G, which were analyzed using a Chi-square test. (*P < 0.05; **P < 0.01; ***P < 0.001).

### MERC disruption paradoxically enhances SOCE via STIM1 activation

Having established that MERCs drive spontaneous mitochondrial Ca^2+^ flickers, we next investigated the functional consequences of this activity for global Ca^2+^ homeostasis. Specifically, we examined the impact of MERC disruption on SOCE—the primary mechanism by which cells replenish ER Ca^2+^ stores. Because MERC-mediated Ca^2+^ transfer continuously drains local ER Ca^2+^, we initially predicted that eliminating MERCs would reduce ER depletion and thus attenuate SOCE. Contrary to this expectation, structural uncoupling via shMFN2, shPDZD8 or shFUNDC1 did not reduce SOCE and even increased thapsigargin-induced ER Ca^2+^ leakage (**Figure S3A–S3C**). We further utilized genetically encoded indicators GCaMPer and T1ER, confirming that MERC disruption elevated resting ER Ca^2+^ levels (**Figure S3D** and **S3E**). Since ER Ca^2+^ stores are primarily maintained by SOCE-mediated refilling, these results suggest that SOCE is actually enhanced by MERC disruption—also hinting that a compensatory mechanism masks this underlying hyperactivation under basal conditions.

We reasoned that microtubule (MT)–ER contacts, which have been shown to sequester STIM1 via the plus-end tracking protein EB1 (Chang *et al*., 2018) were a plausible candidate for this masking effect. To test whether MT-mediated STIM1 sequestration was concealing an underlying SOCE phenotype, we pre-treated cells with the MT-depolymerizing agent colchicine. Under these conditions, MERC disruption (shMFN2, shPDZD8, or shFUNDC1) produced a massive enhancement of SOCE peak amplitude while TG- or histamine-induced ER Ca^2+^ release remained consistently increased (**Figure 3A–3C** and **S3G**). Importantly, the accompanying elevation of intra-ER Ca^2+^ and hyperactive SOCE were both completely abolished by simultaneous knockdown of STIM1 and ORAI1 (**Figure 3F** and **S3F**), confirming that the phenotype was mediated through the canonical SOCE machinery. These results reveal that MERCs do not promote SOCE as initially anticipated; rather, they primarily function to suppress it.

**Figure 3.**
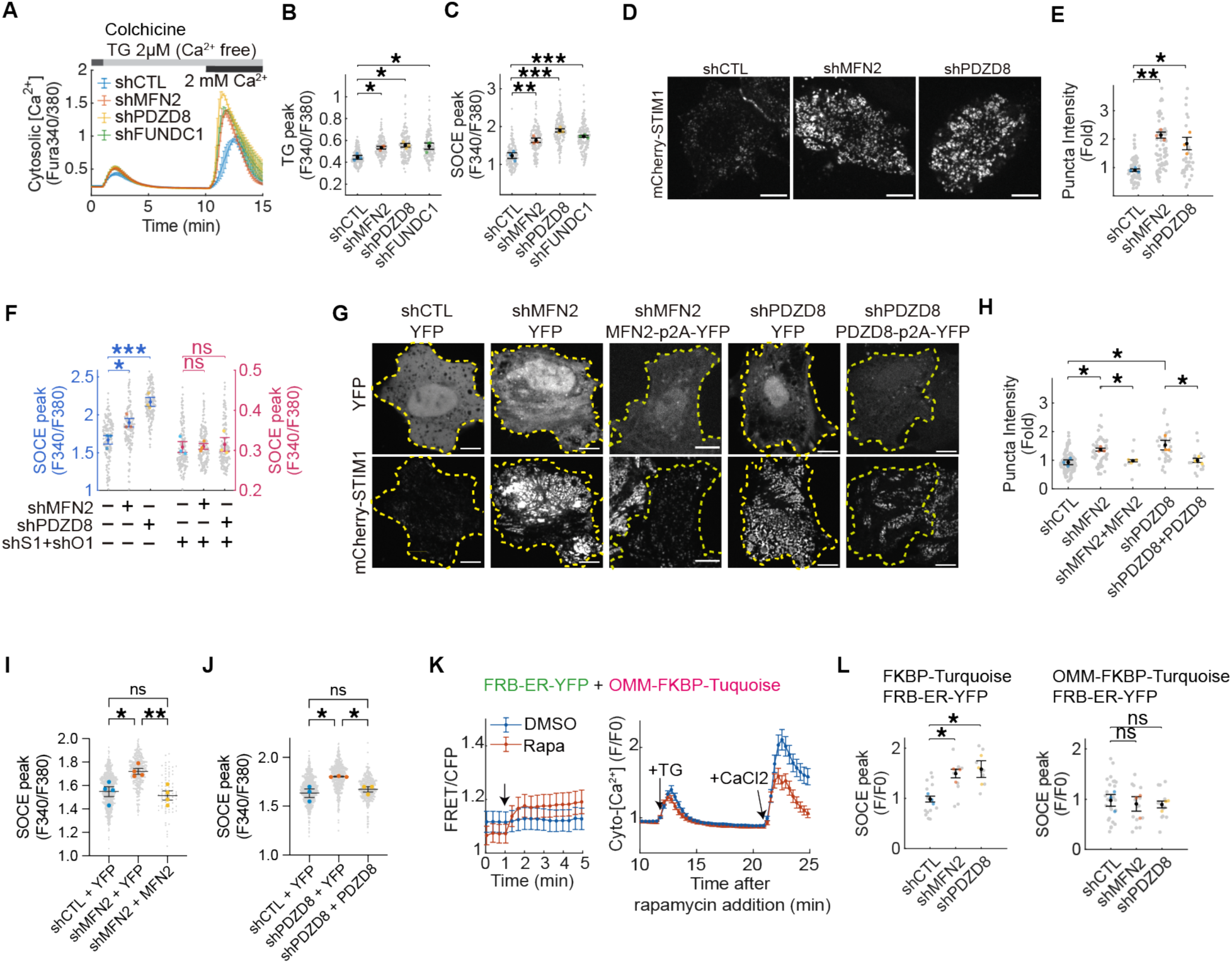
MERC disruption paradoxically enhances SOCE via STIM1 activation. **A.** Cytosolic Ca^2+^ measurements using Fura-2 in control (shCTL) and knockdown (shMFN2, shPDZD8, shFUNDC1) HeLa cells following treatment with 2 µM thapsigargin (TG) in Ca^2+^-free medium to deplete ER stores, followed by the addition of 2mM CaCl_2_ to trigger SOCE. **B–C.** Quantification of the TG-induced ER Ca^2+^ release peak (**B**) and the subsequent SOCE peak (**C**) demonstrates increased Ca^2+^ storage and enhanced SOCE in MFN2-, PDZD8-, and FUNDC1- knockdown cells. **D–E.** MERC disruption promotes STIM1 accumulation at ER-PM junctions. Representative images of mCherry-STIM1 puncta formation after TG treatment in the indicated knockdown cells (**D**), and corresponding quantification of STIM1 puncta intensity (**E**). Scale bar, 10µm. **F.** The hyperactive Ca^2+^ influx is STIM1/ORAI1-dependent. Comparison of SOCE in shMFN2- or shPDZD8-treated cells without (left, blue) or with (right, red) concurrent knockdown of STIM1 and ORAI1 (shS1+shO1). The abrogation of the hyperactive Ca^2+^ influx by shS1+shO1 confirms that MERC disruption promotes Ca^2+^ influx specifically via the STIM1/ORAI1-dependent SOCE pathway. **G–J.** Genetic reconstitution of MERC tethers reverses the hyperactive phenotypes. (**G**) Representative images showing that re-expression of MFN2-p2A-YFP and PDZD8-p2A-YFP in their respective knockdown cells restores STIM1 puncta back to baseline levels. (**H–J**) Quantification confirming that genetic rescue of MFN2 or PDZD8 concurrently normalizes the STIM1 puncta intensity (**H**) and the SOCE peaks (**I, J**). Statistical analysis was performed using one-way ANOVA followed by Tukey’s multiple comparisons test. **K.** Induction of synthetic MERCs specifically suppresses SOCE hyperactivation. (Left) FRET/CFP ratio monitoring the rapid formation of rapamycin-induced contacts in cells co-expressing FRB-ER-YFP and OMM-FKBP-Turquoise. (Right) Representative cytosolic Ca^2+^ traces (measured using R-GECO1) showing the SOCE response in HeLa cells following rapamycin (+Rapa) or vehicle (DMSO) treatment. **L.** Induction of mitochondria-ER tethering rescues SOCE hyperactivation in MERC-deficient cells. Quantification of SOCE following rapamycin-induced formation of either non-specific contacts (using FRB-ER-YFP and cytosolic FKBP-Turquoise) or specific MERCs (using FRB-ER-YFP and OMM-FKBP-Turquoise) in knockdown cells. Data are normalized to the shCTL group. Quantitative data are presented as swarm charts with mean ± SEM. Small gray dots represent individual cells, while larger colored dots indicate the means of at least three independent replicates. Unless otherwise indicated, all statistical analysis was performed using one-way ANOVA followed by Dunnett’s multiple comparisons test. *p < 0.05, **p < 0.01, ***p < 0.001.

To understand the structural basis of this enhanced SOCE, we examined the architecture of ER-PM junctions — the physical platforms for STIM1–ORAI1 complex formation. Strikingly, MERC disruption significantly expanded the overall coverage of ER-PM junctions (**Figure S3H**), indicating the loss of mitochondrial tethering leads to spatial redistribution of the ER membrane toward the cell cortex. Consistent with this structural remodeling, MERC-deficient cells showed increased STIM1 oligomerization and puncta formation (**Figure 3D, 3E, S3I** and **S3J**), as well as enhanced STIM1-ORAI1 colocalization (**Figure S3K** and **S3L**), together linking the expanded ER-PM junctions to the functional Ca^2+^ hyperactivation.

To establish causality between MERC disruption and SOCE hyperactivation, we performed two orthogonal rescue experiments. Genetic reconstitution of MFN2 or PDZD8 in knockdown cells reversed both STIM1 puncta (**Figure 3G** and **3H**) and SOCE enhancement (**Figure 3I** and **3J**). Crucially, since these assays were conducted under full thapsigargin-induced store depletion, genetic re-expression of MFN2 or PDZD8 normalized STIM1 puncta intensity by restoring the structural trapping and spatial competition at MERCs rather than altering baseline ER fill state. In parallel, acute rapamycin-induced MERC formation in MERC-deficient cells suppressed SOCE (**Figure 3K** and **S3M**) and abolished the hyperactivation phenotype (**Figure 3L** and **S3N**).

Ultimately, these findings demonstrate that re-establishing physical ER-mitochondria proximity restores normal SOCE—independent of the specific tethering protein used—confirming that the suppressive function of MERCs resides in the spatial coupling itself, rather than in the distinct signaling properties of individual tethers.

### MERCs suppress SOCE via IP₃R–VDAC1-mediated ER–mitochondrial Ca^2+^ transfer

To delineate the molecular mechanism underlying MERC-mediated SOCE suppression, we first confirmed that the phenotype was not a consequence of altered protein abundance: RT-qPCR and Western blots revealed that mRNA and protein levels of STIM1 and ORAI1 remained unchanged in MFN2- or PDZD8-knockdown cells (**Figure S4A, S4B, S4C** and **S4D**). Moreover, the hyperactive SOCE could be fully rescued by re-establishing MERCs via the rapamycin-inducible system (**Figure 3L**), indicating that the regulatory effect depends on ER-mitochondrial spatial proximity rather than specific tethering proteins. We thus adopted an exhaustive approach to systematically evaluate three candidate mechanisms underlying MERC-mediated SOCE suppression (**Figure 4A**): (i) modulation by mitochondrial metabolic outputs or ROS; (ii) regulation of STIM1 activities by STIM1 regulators; or (iii) ER-to-mitochondria Ca^2+^ transfer-induced changes in STIM1 dynamics.

**Figure 4.**
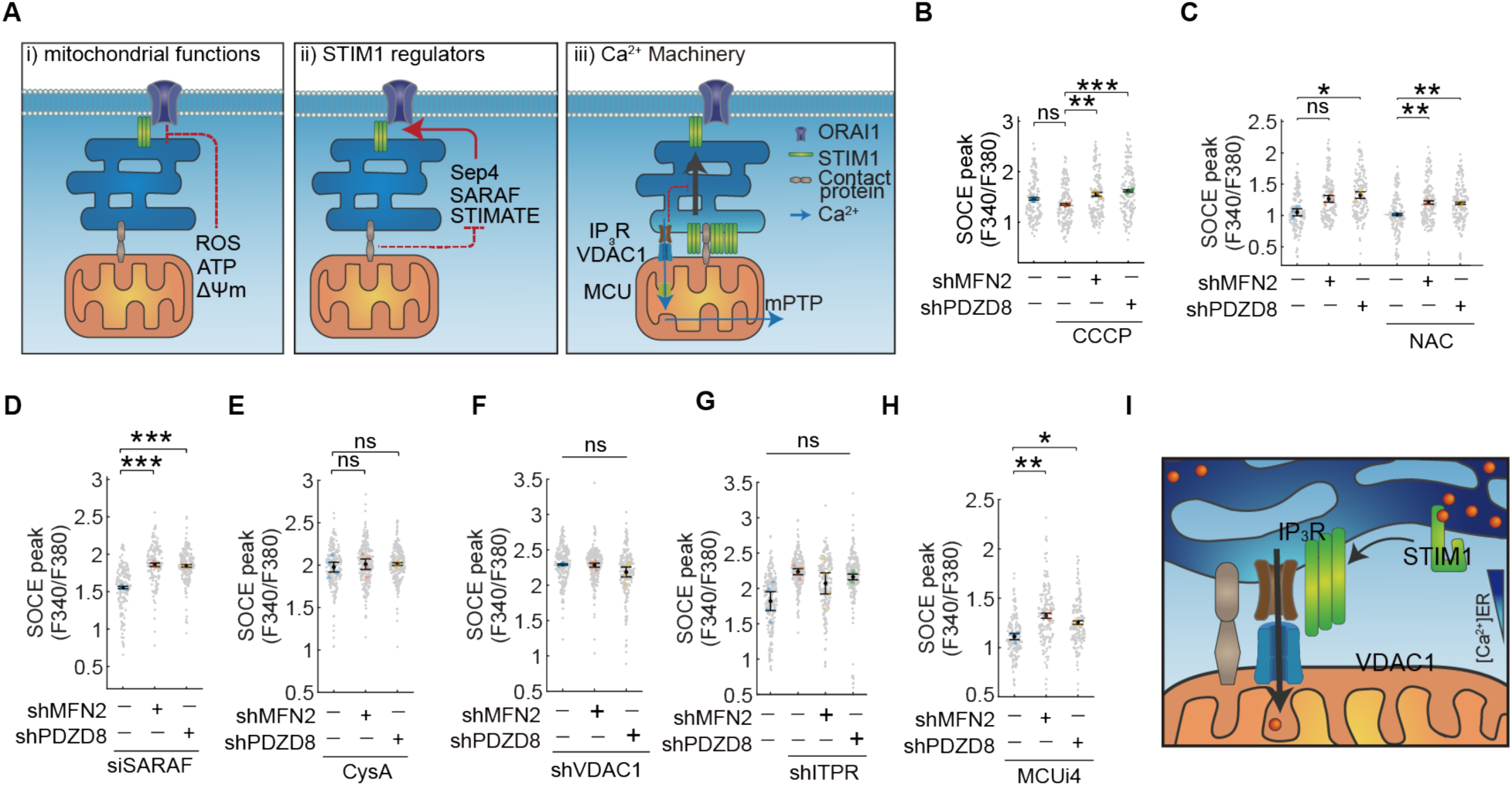
MERCs suppress SOCE via IP₃R–VDAC1-mediated ER–mitochondrial Ca^2+^ transfer, independent of mitochondrial functions or STIM1 regulators. **A.** Schematic representation of three proposed hypotheses tested for the underlying mechanisms of SOCE suppression at MERCs: (i) modulation by mitochondrial metabolic outputs or reactive oxygen species (ROS); (ii) STIM1 regulators including SARAF, septins and STIMATE; and (iii) mitochondrial Ca^2+^ flux through IP_3_R, MCU and mPTP. **B–C.** Evaluation of hypothesis (i). Quantification of SOCE peak amplitudes reveals that neither the disruption of mitochondrial membrane potential (ΔΨm) by CCCP (**B**) nor the scavenging of ROS by N-acetylcysteine (NAC) (**C**) abolishes the SOCE hyperactivation induced by MERC depletion (shMFN2 or shPDZD8). **D.** Evaluation of hypothesis (ii). Genetic knockdown of the STIM1 regulatory protein SARAF fails to prevent SOCE upregulation in MERC-deficient cells. **E–H.** Evaluation of hypothesis (iii). (**E**) Cyclosporin A (CysA) treatment completely abolishes SOCE hyperactivation in MERC-deficient cells. (**F**) Knockdown of the outer mitochondrial membrane channel VDAC1 eliminates SOCE hyperactivation in MERC-deficient cells. (**G**) Knockdown of the ER Ca^2+^ release channel IP_3_R similarly eliminates SOCE hyperactivation in MERC-depleted cells. (**H**) Inhibition of MCU with MCUi4 fails to abolish the hyperactive SOCE in MERC-deficient cells. Collectively, these findings indicate that MERCs suppress SOCE by facilitating the local transfer of Ca^2+^ from the ER into the mitochondrial intermembrane space (IMS), rather than through Ca^2+^ uptake from IMS into the mitochondrial matrix. **I.** Proposed mechanistic model. Through systematic exploration, the data pinpoint the IP_3_R–VDAC1 Ca^2+^ channeling complex as the indispensable machinery for MERC-regulated SOCE. This supports the hypothesis that active Ca^2+^ transfer at MERCs creates a localized low-Ca^2+^ microdomain at the ER side of MERCs to trap STIM1, thereby sequestering it from the plasma membrane and preventing SOCE overactivation. Cells were pretreated with 10 μM colchicine to prevent potential interference of microtubules. Quantitative data are presented as swarm charts with mean ± SEM. Small gray dots represent measurements from individual cells, while larger colored dots indicate the means of at least three independent biological replicates. Statistical analysis was performed using one-way ANOVA followed by Dunnett’s multiple comparisons test. *p < 0.05, **p < 0.01, ***p < 0.001.

To test hypothesis (i), we disrupted mitochondrial membrane potential (ΔΨm) using CCCP or FCCP; neither intervention abolished the hyperactive SOCE in MERC-deficient cells (**Figure 4B** and **S4E**), ruling out ΔΨm-dependent mechanisms. Consistently, inhibition of ATP synthase with oligomycin A, or of the electron transport chain with rotenone and antimycin A, likewise had no effect (**Figure S4F** and **S4G).** Furthermore, while baseline ROS levels showed no significant changes in MERC-deficient cells (**Figure S4H** and **S4I**), neutralizing ROS with either the general scavenger NAC (**Figure 4C**) or the mitochondria-targeted antioxidant mitoTEMPO (**Figure S4J**) failed to attenuate the hyperactive SOCE in MERC-deficient cells. Together, these results exclude mitochondrial metabolic outputs and oxidative stress as primary drivers.

Regarding hypothesis (ii), we depleted a panel of known STIM1 regulators to determine if they mediated MERC-mediated SOCE suppression. Knockdown of SARAF (Palty *et al*., 2012) (**Figure 4D** and **S4K**) failed to abolish the hyperactive SOCE in MERC-deficient cells. Similarly, knocking down STIMATE (Jing *et al*., 2015) or septins including septin-4 and septin-7 (Deb & Hasan, 2019; Deb *et al*, 2016; Katz *et al*, 2019; Sharma *et al*, 2013; Yang *et al*, 2010)) (**Figure S4L–S4O**) produced no rescue. The consistent failure of these interventions rules out canonical STIM1 regulatory pathways as the primary mechanism underlying MERC-mediated SOCE suppression.

Having excluded metabolic and canonical regulatory mechanisms, we turned to hypothesis (iii) by focusing on the inter-organelle Ca²⁺ transfer machinery itself. Notably, cyclosporin A (CysA), which inhibits cyclophilin D-mediated mitochondrial permeability transition pore (mPTP) opening, eliminated the effects of MERC disruption on SOCE (**Figure 4E**). Since mPTP activity triggers mitochondrial ΔΨm collapse, ROS generation and Ca^2+^ transfer—each of which conceivably modulates SOCE through independent pathways—we interrogated whether these downstream consequences accounted for the CysA effect. Notably, neither CCCP / FCCP-mediated dissipation of the mitochondrial ΔΨm (**Figure 4B** and **S4E**), nor NAC / mitoTEMPO-mediated ROS scavenging (**Figure 4C**) phenocopied CysA-mediated SOCE rescue. Since VDAC1 is a key regulatory component of mPTP complex (Javadov *et al*, 2009), we investigated the role of VDAC1-mediated Ca^2+^ transfer at MERCs. Indeed, concurrent depletion of either *VDAC1* (shVDAC1; **Figure 4F**) or its ER counterpart *IP_3_R* (shITPR; **Figure 4G**) in MERC-deficient cells extinguished SOCE hyperactivation. This epistatic rescue indicates that MERC-mediated SOCE regulation requires the IP_3_R-VDAC1 axis, a finding further corroborated by the observation that knockdown of IP_3_R or VDAC1 alone was sufficient to trigger SOCE hyperactivation, effectively phenocopying the loss of physical MERC tethers (**Figure S4P**).

In contrast, inhibition of the MCU using MCUi4 (Di Marco *et al*, 2020), DS16570511 (Kon *et al*, 2017), or ruthenium red (Moore, 1971) failed to abolish hyperactive SOCE (**Figure 4H** and **S4Q, R**). MCU inhibition also did not reverse the elevated steady-state ER Ca^2+^ in MERC-deficient cells (**Figure S4S, T**). NCLX expression likewise remained unchanged (**Figure S4U**). These data seamlessly reconcile the apparent paradox with our earlier observation that, while MCU activity is strictly required for matrix-targeted biosensors to visually register the flickering transients (**Figure 1L, M**), the actual physical clearance of Ca²⁺ out of the ER lumen occurs at the intermembrane space (IMS) interface via the IP₃R–VDAC1 complex.

Collectively, these findings indicate that MERCs suppress SOCE by facilitating the local transfer of Ca^2+^ from the ER into the mitochondrial intermembrane space (IMS) via the IP_3_R–VDAC1 complex, rather than through Ca^2+^ uptake into the mitochondrial matrix (Javadov *et al*., 2009). Through this systematic exploration, we confirm that the IP_3_R-VDAC1 complex at the MERCs acts as the indispensable core machinery restricting SOCE. Based on these results, we propose a mechanistic model where IP_3_R–VDAC1-mediated Ca^2+^ transfer at MERCs creates a localized low-Ca^2+^ microdomain at the ER side of the junction to trap STIM1, thereby sequestering it from the plasma membrane and preventing SOCE overactivation (**Figure 4I**).

### MERCs reduce SOCE by trapping STIM1 through local ER Ca^2+^ depletion and competitive sequestration

Having identified IP₃R-VDAC1-mediated Ca²⁺ transfer as the essential trigger, we next investigated the downstream mechanism within the ER that couples this localized Ca²⁺ drain to suppressed SOCE. Since IP₃R-VDAC1 complexes continuously siphon Ca²⁺ from the peri-MERC ER lumen, we hypothesized that this creates a sustained local Ca²⁺ depletion microdomain sufficient to activate STIM1 specifically at MERCs. This would sequester STIM1 away from ER-PM junctions, thereby suppressing SOCE (**Figure 4I**). In this model, MERCs function as structural traps for STIM1. Their disruption would consequently release a larger STIM1 pool to drive hyperactive Ca²⁺ entry.

To directly test this hypothesis, we acutely induced synthetic MERCs using a rapamycin-inducible system while simultaneously monitoring real-time ER Ca²⁺ and STIM1 dynamics. Using the ER-targeted Ca^2+^ indicator ER-LAR-GECO1, we observed a rapid and spatially restricted decrease in ER Ca^2+^ precisely at the newly formed contact sites (**Figure 5A-C** and **S5A-C**). Kymograph analyses confirmed that this contact formation was rapid and spatially restricted to the induced MERCs (**Figure S5**). Concomitantly, mCherry-STIM1 was rapidly recruited to and enriched at these synthetic MERCs (**Figure 5D-F**). These results provide direct visual evidence that local Ca^2+^ depletion at MERCs is sufficient to sequester STIM1 in situ.

**Figure 5.**
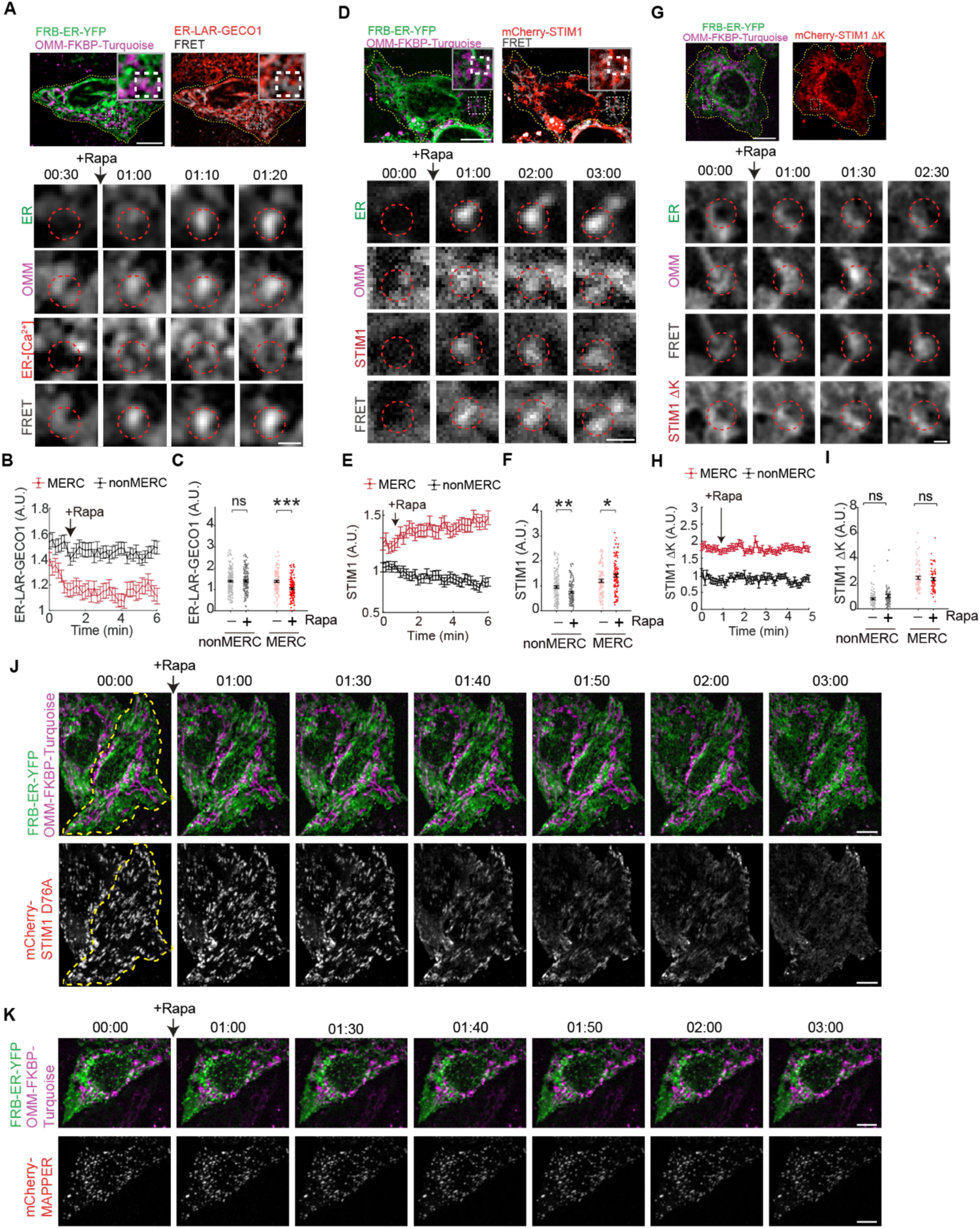
MERCs reduce SOCE by trapping STIM1 through local ER Ca^2+^ depletion and competitive sequestration. **A–C**. Acute MERC induction triggers localized ER Ca^2+^ depletion. Cells co-expressing the rapamycin-inducible tethering system (FRB-ER-YFP and OMM-FKBP-Turquoise) and an ER-targeted Ca^2+^ sensor (ER-LAR-GECO1) were imaged via time-lapse microscopy. Addition of rapamycin (Rapa) triggers a rapid and localized decrease in ER Ca^2+^ specifically at the newly formed MERCs, as shown in representative images (**A**) and normalized intensity quantification (**B, C**). **D–F**. STIM1 is rapidly sequestered at induced MERCs. Upon rapamycin addition, wild-type mCherry-STIM1 is rapidly recruited to and enriched at synthetic contact sites. Representative images (**D**) and quantification (**E, F**) demonstrate the localized accumulation of STIM1 following MERC formation. **G–I.** The STIM1 polybasic domain is required for localized trapping at MERCs. Unlike the wild-type protein, the STIM1 ΔK mutant, which lacks the polybasic domain (residues 671-685), fails to show localized enrichment at induced MERCs. Representative images (**G**) and quantification (**H, I**) reveal that the spatial sequestration of STIM1 at MERCs depends on its C-terminal polybasic motifs. **J.** MERC induction recruits pre-activated STIM1 from ER-PM junctions. Representative images of the constitutively active STIM1 D76A mutant. Acute MERC induction disassembles pre-existing STIM1 D76A puncta at ER–PM junctions by relocating the STIM1 protein toward MERCs. **K.** MERC induction does not compromise the structural integrity of ER–PM junctions. Representative time-lapse images showing persistent mCherry-MAPPER puncta (marking ER–PM junctions) during acute MERC induction. Unlike the displacement of pre-activated STIM1 seen in (**J**), the stability of MAPPER signals confirms that STIM1 redistribution is driven by competitive sequestration at MERCs rather than physical disassembly of ER–PM junctions. Insets (white dashed boxes) show magnified regions of interest. Red dashed circles indicate sites of MERC formation following rapamycin induction. Data are represented as mean ± SEM. Statistical significance was determined by the Mann-Whitney U test (**p<0.01, ***p<0.001). Scale bar: 10 μm; inset, 2 μm.

To dissect the structural requirements for this sequestration, we utilized two well-characterized STIM1 mutants. First, we examined the STIM1 ΔK mutant, which lacks the C-terminal polybasic tail (residues 671–685; hereafter referred to as the “polybasic K-domain”) required for electrostatic membrane anchoring (Liou *et al*, 2007; Yuan *et al*, 2009). While STIM1 ΔK retains normal luminal Ca^2+^ sensing and oligomerization capabilities, it completely failed to accumulate at the rapamycin-induced MERCs (**Figure 5G–I**). This result indicates that local Ca^2+^ depletion alone is insufficient for STIM1 trapping. The polybasic K-domain is required for STIM1 to be sequestered within the MERC microenvironment. Complementarily, we utilized the constitutively active STIM1 D76A mutant, which mimics the store-depleted state (Liou *et al*., 2005; Zhang *et al*, 2005). Upon MERC induction, pre-existing STIM1 D76A puncta at ER-PM junctions were actively disassembled, probably redistributed toward the newly formed MERCs (**Figure 5J**). Crucially, this spatial redistribution was not due to a global collapse of the ER architecture, as the ER-PM junction marker MAPPER remained relatively stable throughout acute MERC induction (**Figure 5K**). Together, these data establish that IP₃R-VDAC1-mediated Ca²⁺ transfer at MERCs generates a sustained local ER Ca²⁺ depletion microdomain that engages STIM1 via its polybasic K-domain. This process sequesters STIM1 away from the plasma membrane through competitive recruitment, thereby suppressing SOCE.

### A tripartite PM-mitochondria-microtubule system regulates SOCE through competitive STIM1 sequestration

Our data so far has established MERCs as the primary checkpoint for STIM1 sequestration. However, a puzzling observation remained: under basal conditions without MT disruption, MERC disruption alone did not significantly alter SOCE amplitude, suggesting that a compensatory mechanism was buffering the excess STIM1 released from disrupted MERCs. The MT plus-end tracking protein EB1 has been shown to bind and restrain STIM1 at growing MT tips, restricting its translocation to the plasma membrane (Chang *et al*., 2018; Grigoriev *et al*, 2008; Honnappa *et al*, 2009).

We therefore hypothesized that upon MERC disruption, the MT-EB1 comets intercept the liberated STIM1 pool and serve as a secondary safeguard against SOCE over-activation. Building on this reasoning and our earlier functional findings, we conceptualized a tripartite spatial competition model in which STIM1 distribution is dynamically negotiated among three structural hubs: MERCs, ER-PM junctions, and MT-EB1 comets (**Figure 6A**).

**Figure 6.**
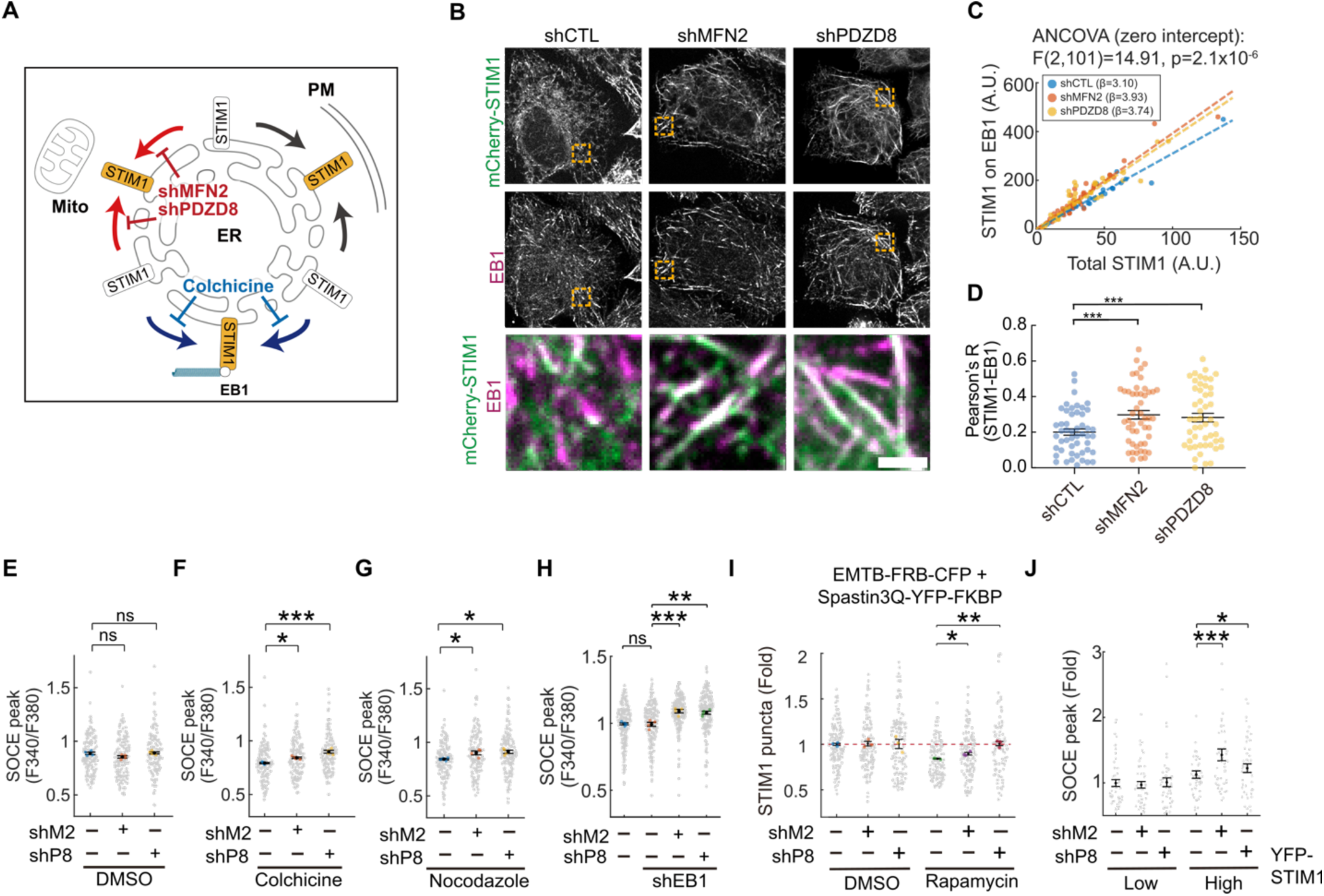
A tripartite PM-mitochondria-microtubule system regulates SOCE through competitive STIM1 sequestration. **A.** Working hypothesis of the tripartite spatial balance. Schematic illustrating the dynamic competition for STIM1 distribution among three structural hubs: MERCs, ER-PM junctions, and the MT (EB1)-ER contacts. **B–D.** Visualization and quantification of the STIM1-EB1 interaction. (**B**) Representative images of STIM1 (mCherry-STIM1, green) and endogenous EB1 comets (EB1, magenta) in control (shCTL) and MERC-disrupted (shMFN2 and shPDZD8) HeLa cells. Orange boxes indicate regions magnified in the bottom panels. Scale bar: 1 μm. (**C**) ANCOVA (zero intercept) of STIM1 intensity on EB1 versus total STIM1; the slope (β) reflects STIM1 recruitment efficiency to EB1. Both shMFN2 (β = 3.93) and shPDZD8 (β = 3.74) show steeper slopes than shCTL (β = 3.10), F(2,101) = 14.91, p = 2.1 × 10⁻⁶. (**D**) Per-cell Pearson’s correlation between STIM1 and EB1 signals. **E–I.** Impact of microtubule disruption on SOCE and STIM1 translocation. (**E–G**) Quantification of SOCE peak amplitudes (F340/F380) in shCTL, shMFN2, and shPDZD8 HepG2 cells following global microtubule (MT) depolymerization with DMSO (vehicle) (**E**) or colchicine (**F**) or nocodazole (**G**). (**H**) SOCE peak amplitudes in the indicated knockdown cells upon genetic depletion of EB1 (shEB1). (**I**) Quantification of STIM1 puncta intensity following rapamycin-induced MT disassembly using EMTB-FRB-CFP and Spastin3Q-YFP-FKBP system, demonstrating that MT disassembly unmasks the significant accumulation of STIM1 puncta in MERC-deficient cells. **J.** Finite sequestration capacity of the tripartite system. Overexpression of STIM1 in MERC-deficient cells overwhelms the MT-EB1 buffer and produces massive SOCE hyperactivation, demonstrating that the combined sequestration capacity of MERCs and MTs is finite. Quantitative data are presented as swarm charts with mean ± SEM. Small gray dots represent measurements from individual cells, while larger colored dots indicate the means of at least three independent replicates. Statistical analysis was performed using one-way ANOVA followed by Dunnett’s multiple comparisons test. *p < 0.05, **p < 0.01, ***p < 0.001.

A key prediction of this model is that MERC disruption should increase STIM1 loading onto EB1-labeled MT plus-ends. To test this directly, we quantified STIM1–EB1 colocalization by confocal imaging in shCTL, shMFN2, and shPDZD8 cells (**Figure 6B**). To account for cell-to-cell variability in STIM1 expression, we used zero-intercept ANCOVA to compare the relationship between STIM1 intensity on EB1 comets and total cellular STIM1 across the three groups. The regression slope, which reflects STIM1 recruitment efficiency per unit of total STIM1, was significantly higher in both shMFN2 (β = 3.93) and shPDZD8 (β = 3.74) cells than in shCTL (β = 3.10) [F(2,101) = 14.91, p = 2.1 × 10⁻⁶; **Figure 6C**]. Consistently, the per-cell Pearson’s correlation coefficient between STIM1 and EB1 signals was elevated in both knockdown conditions compared with control (**Figure 6D**), indicating increased spatial overlap of the two proteins. Together, these results demonstrate that loss of MERC integrity enhances STIM1 recruitment to growing MT plus-ends.

To establish the functional significance of this secondary buffer, we employed three orthogonal approaches to disrupt MT-mediated STIM1 sequestration. First, while global MT depolymerization using colchicine or nocodazole produced SOCE hyperactivation in MFN2- and PDZD8-depleted cells (**Figure 6E–G**), it had no significant effect on TG-induced ER Ca^2+^ leakage (**S6A–C**), which represents the basal ER Ca^2+^ store. Because the STIM1 requirement for basal ER homeostasis is much lower than the demand during a maximal SOCE peak, these results indicate that MT-EB1 comets possess a finite STIM1 sequestration capacity—sufficient to regulate peak SOCE without compromising the basal ER Ca^2+^ store. Notably, MT stabilization with paclitaxel did not alter the effects of MERC-depletion on TG-induced ER Ca^2+^ leakage and SOCE activation (**Figure S6D, E**), suggesting that the buffering capacity depends on the availability of MT-based sequestration sites rather than MT dynamics. Second, genetic depletion of EB1 (shEB1) recapitulated this SOCE enhancement in MERC-deficient cells (**Figure 6H**), confirming that the buffering function operates specifically through the STIM1-EB1 interaction rather than through nonspecific alterations of cellular architecture. Third, acute rapamycin-inducible Spastin (Liu *et al*, 2022) (**Figure S6F, G**) exacerbated STIM1 puncta formation in MFN2- or PDZD8-depleted cells (**Figure 6I** and **S6H**). This rapid induction demonstrates that the hyperactivation phenotype is a direct consequence of MT loss rather than an artifact of prolonged drug exposure. Together, these three orthogonal interventions consistently demonstrate that the MT-EB1 comets function as a genuine secondary safeguard that counteracts MERC loss by sequestering surplus STIM1.

We further explored whether the microtubule-dependent modulation of SOCE involved direct structural changes to organelle architecture. Using 3D volumetric reconstruction, we found that microtubule depolymerization via colchicine treatment significantly increased the physical coupling between the ER and mitochondria, resulting in a marked expansion of both ER-contacted mitochondrial surface and mitochondria-contacted ER surface (**Figure S6I**). This architectural remodeling reveals that microtubules normally restrict the available ER surface area for mitochondria–ER tethering. Consequently, the expansion of the mitochondrial hub in microtubule-deficient cells creates a larger “MERC trap” that enhances the sequestration of STIM1 within local ER Ca^2+^ depletion microdomains. This structural expansion provides a mechanistic explanation for the reduction in peak SOCE observed in control cells following microtubule disruption by colchicine, nocodazole, or acute Spastin3Q-mediated disassembly (**Figure 6F, G, I**).

Finally, we investigated whether this secondary buffer operates with a finite capacity. Two stoichiometric experiments addressed this question. Overexpression of STIM1 in MERC-deficient cells overwhelmed the MT-EB1 buffer and produced massive SOCE hyperactivation (**Figure 6J**), demonstrating that when the total STIM1 pool exceeds the combined sequestration capacity of MERCs and MTs, the surplus STIM1 inevitably reaches the plasma membrane. Conversely, EB1 overexpression reduced SOCE in control cells as expected, but failed to rescue the elevated SOCE in MERC-deficient cells (**Figure S6J**), revealing that when the primary MERC trap is absent, the quantity of liberated STIM1 exceeds what even an augmented MT-EB1 sink can accommodate. Together, these findings reveal an exquisite stoichiometric balance: when MERCs alone are disrupted, the MT-EB1 comets actively intercept the released STIM1 pool to maintain SOCE; however, their simultaneous failure precipitates pathological Ca²⁺ overload. Collectively, our data uncover an integrated tripartite regulatory network in which ER, mitochondria, and MT (EB1) cooperate through competitive STIM1 sequestration to maintain Ca²⁺ homeostasis (**Figure 7**).

**Figure 7.**
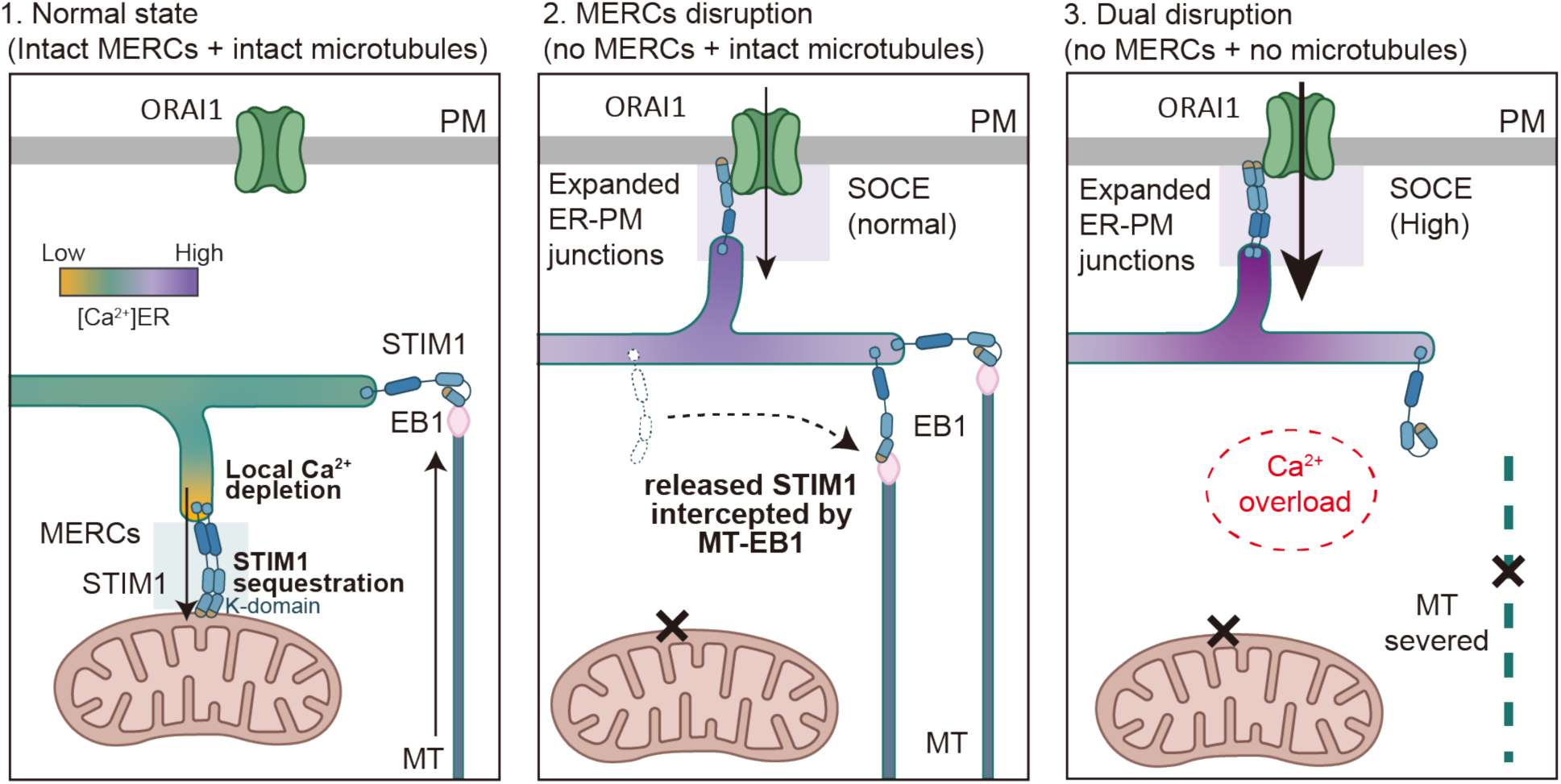
Model of the integrated tripartite STIM1 regulatory network. (1) Normal state. STIM1 is partitioned between MERCs and the MT (EB1)-ER contacts, which collectively prevent excessive STIM1 translocation to the PM and maintain baseline SOCE. (2) MERC disruption. STIM1 released from the mitochondrial hub is intercepted by the MT (EB1)-ER contacts, preventing SOCE overactivation despite the presence of expanded ER-PM junctions. (3) Dual disruption. Without both sequestration hubs, unrestrained STIM1 floods the ER-PM junctions, leading to Ca^2+^ overload.

## DISCUSSION

Our study establishes that MERCs function as primary spatial checkpoints of SOCE by generating localized ER Ca²⁺ depletion microdomains that competitively sequester STIM1. This work advances the understanding of mitochondrial Ca²⁺ dynamics by directly linking stochastic organelle-level events to global (cellular level) Ca²⁺ homeostasis through a defined molecular mechanism. While mitochondrial miniature Ca^2+^ signals driven by ryanodine receptors had been noted (Pacher *et al*., 2002), subsequent work identified ER-to-mitochondria Ca^2+^ flux at MERCs as a key driver of local mitochondrial Ca²⁺ events (Cardenas *et al*., 2020; Cardenas *et al*., 2010; Csordas *et al*., 2006; Csordas *et al*., 2010; Rizzuto *et al*., 1998). Our study further advances the field in two respects. First, we provide a comprehensive quantitative characterization of these events under resting physiological conditions—defining their frequency (∼1.5 min⁻¹ per mitochondrion), amplitude (∼0.9–1.3 fold of baseline), and duration (6–33 seconds)—and demonstrating that they follow a Poisson distribution. This statistical profile establishes that each mitochondrion operates as an autonomous stochastic unit, capable of independent spatiotemporal regulation of local Ca²⁺ dynamics. Second, we identify the functional consequence of these events: MERC-mediated flickering siphons local ER Ca²⁺ to trap STIM1, serving as a previously unrecognized safeguard against Ca²⁺ overload. Critically, this MERC-based STIM1 trapping requires the participation of the PM and MTs. Loss of MERCs enhances ER-PM expansion for SOCE hyperactivation, while MT-EB1 comets serve as a finite-capacity checkpoint for STIM1 sequestration when the primary MERC trap is compromised. Together, these findings define an integrated tripartite regulatory network in which the ER, mitochondria, and the microtubule cytoskeleton cooperate to maintain Ca²⁺ homeostasis.

We further demonstrate that these flickers are MCU-dependent yet entirely independent of ROS signaling. While mitochondrial ROS and Ca²⁺ were proposed to be co-regulated (Booth *et al*., 2021; Patel *et al*., 2022) and the concept of superoxide flashes (Wang *et al*, 2008) has been debated, our data clearly dissociate ROS and mitochondrial Ca^2+^ flickering. Simultaneous monitoring with matrix-roGFP confirmed that no spontaneous redox fluctuations coincided with Ca^2+^ flickers (**Figure S1C** and **S1D**). Furthermore, mitoPQ-induced ROS elevation failed to alter Ca^2+^ flickering (**Figure S1E–G**). Similarly, ROS scavenging with NAC or mitoTEMPO also did not change the SOCE phenotype (**Figure 4C** and **S4J**). Together with the sensitivity of Ca^2+^ flickers to MCU inhibition and MERC disruption, this ROS independence confirms the IP_3_R-VDAC1 complex as the primary driver of these events, which is consistent with the principle that MCU activation requires the high-Ca^2+^ microdomains generated at MERCs.

Our findings also extend the current conceptual framework of how mitochondria influence SOCE, from a purely positive role via Ca^2+^ buffering to a suppressive role via STIM1 sequestration. Early studies revealed that mitochondria act as positive modulators of SOCE through two complementary mechanisms: (1) Subplasmalemmal mitochondria buffer Ca²⁺ in the immediate vicinity of CRAC channels, preventing Ca²⁺-dependent inactivation and sustaining Ca²⁺ influx (Gilabert & Parekh, 2000; Hoth *et al*, 1997; Malli *et al*, 2003). (2) Mitochondrial Ca^2+^ uptake from the ER at MERCs reduces ER Ca^2+^ store, thereby prolonging the store-depleted state that maintains STIM1 activation (Csordas *et al*., 2010; Deak *et al*, 2014). In both cases, mitochondria enhance the duration of SOCE activity via Ca^2+^ buffering. In contrast, our study reveals a distinct mechanism wherein MERCs suppress the magnitude of SOCE activity via STIM1 sequestration. This counterintuitive conclusion began with the known phenomenon that MERC elimination increased rather than decreased intra-ER Ca²⁺ storage. While previous studies primarily attributed elevated ER Ca^2+^ in MERC-deficient cells to reduced ER-to-mitochondria Ca²⁺ transfer (Csordas *et al*., 2010; Lee *et al*, 2018), our data argue that this elevation is instead a downstream consequence of SOCE hyperactivation, since simultaneous STIM1/ORAI1 knockdown abolished the phenotype (**Figure S3F**). This finding indicates that enhanced SOCE is the proximate cause of elevated ER Ca^2+^ in MERC-deficient cells, consistent with the previous observation that MFN2 knockdown enhances SOCE (Singaravelu *et al*, 2011). Furthermore, this suppressive mechanism operates independently of mitochondrial Ca²⁺ buffering capacity, as dissipation of the mitochondrial membrane potential (ΔΨm) by CCCP or FCCP failed to reverse the hyperactive SOCE observed in MERC-deficient cells (**Figure 4B** and **S4E**). Together, our findings suggest that mitochondria exert dual controls over SOCE, by promoting SOCE duration through Ca²⁺ buffering, while simultaneously restraining SOCE magnitude through MERC-mediated STIM1 sequestration. The balance between these forces likely serves as a dynamic rheostat to calibrate SOCE according to cellular Ca²⁺ demand.

Our results also advance the current understanding regarding the mechanism of STIM1 activation. It is well established that depletion of intra-ER Ca^2+^ triggers a STIM1 conformational change, initiating its oligomerization and translocation to ER-PM junctions to activate ORAI1-mediated Ca^2+^ influx (Liou *et al*., 2005; Prakriya *et al*., 2006; Roos *et al*., 2005; Stathopulos *et al*, 2013; Vig *et al*., 2006). We further demonstrate that this canonical activation mechanism can be engaged locally at MERCs: by generating mitochondrial Ca^2+^ flickers, IP_3_R-VDAC1-mediated Ca^2+^ transfer simultaneously depletes local ER Ca^2+^ at MERCs, leading to STIM1 accumulation at the contact sites.

The restricted diffusion of Ca^2+^ within the ER lumen (Crapart *et al*, 2024; Csordas *et al*., 2006; Marchi *et al*, 2018) enables continuous Ca^2+^ transfer at MERCs to establish a sustained local Ca^2+^ depletion microdomain. Consistent with this, time-lapse monitoring of ER Ca^2+^ following rapamycin-induced MERC formation revealed a selective and persistent decrease specifically at mitochondria-contacted ER regions, while Ca^2+^ levels at non-contact regions remained stable (**Figure 5A-C**). This sustained local depletion amounted to approximately 20% below baseline (from 1.4 A.U. (100%) to 1.1 A.U. (79%), **Figure 5C**). Notably, the EF-SAM region of STIM1 binds Ca^2+^ with low affinity (K_D_ ∼0.2 to 0.6 mM) (Stathopulos *et al*, 2006; Zheng *et al*, 2008), a range comparable to basal ER luminal Ca²⁺ concentrations (Greotti *et al*, 2016; Rossi & Taylor, 2020). A 20% reduction in local ER Ca²⁺ is therefore sufficient to shift the equilibrium toward STIM1 activation and accumulation at the contact site. This model is complementary to a recent independent study by Orantos-Aguilera et al. (2026), which identified a structural pool of STIM1 residing at MERCs via interaction with GRP75 under resting conditions. In support of this model, our results also demonstrate a substantial constitutive presence of STIM1 at MERCs prior to stimulation (**Figure 5F**). Whereas their work defines the steady-state structural anchoring of STIM1 at MERCs, our findings characterize the activation-state dynamics by which local ER Ca²⁺ depletion microdomains actively recruit and sequester STIM1 in situ—together establishing MERCs as a bona fide spatial anchor for both resting and activated STIM1 pools (Orantos-Aguilera *et al*., 2026).

Our dissection of STIM1 mutants further reveals the structural requirement for STIM1 trapping at MERCs. The STIM1 ΔK mutant, which lacks the C-terminal polybasic domain (residues 671-685) for electrostatic membrane anchoring, fails to accumulate at rapamycin-induced MERCs (**Figure 5G-I**) despite normal Ca^2+^ sensing. This proves that local Ca²⁺ depletion alone is insufficient for STIM1 trapping—physical anchoring via the polybasic K-domain is required to retain activated STIM1 within the MERC microenvironment. The requirement for the K-domain parallels its role in anchoring STIM1 at ER-PM junctions (Liou *et al*., 2007). Recent pull-down data also reveal that this K-domain contributes to STIM1-GRP75 interaction at MERCs (Orantos-Aguilera *et al*., 2026), further supporting the importance of this K-domain for STIM1 localization to MERCs. Furthermore, the constitutively active STIM1 D76A mutant, which localizes to ER-PM junctions even without ER Ca^2+^ depletion, is actively redistributed away from ER-PM junctions under acute MERC induction (**Figure 5J, K**), indicating a dynamic competition for STIM1 between MERCs and ER-PM junctions. Together, these findings establish a dual-layer model of ER-mediated STIM1 regulation: global control at ER-PM junctions through bulk luminal Ca²⁺ depletion, and local control at MERCs through localized Ca²⁺ microdomains.

The observation that cyclosporin A (CysA) abolished SOCE hyperactivation in MERC-deficient cells initially suggested mPTP involvement. CysA inhibits cyclophilin D, thereby suppressing mPTP opening. Since mPTP contributes to physiological mitochondrial Ca²⁺ release, its sustained opening triggers ΔΨm collapse and ROS generation, resulting in SOCE modulation through redox modifications of STIM1, ORAI1 or IP₃R (Booth *et al*., 2021; Patel *et al*., 2022). Moreover, because mPTP opening involves VDAC1 (Javadov *et al*., 2009), it could also result in SOCE modulation via IP_3_R-VDAC1-mediated mitochondrial Ca^2+^ flickering. Therefore, CysA may disrupt MERC-mediated SOCE by altering ΔΨm, ROS and / or VDAC1. However, our pharmacological evidence argued against the involvement of ΔΨm or ROS, as ΔΨm dissipation by CCCP and FCCP (**Figure 4B** and **S4E**) and ROS scavenging by NAC and mitoTEMPO (**Figure 4C** and **S4J**) both failed to phenocopy the CysA effect. This failure indicates that the CysA effect on SOCE does not operate through canonical mPTP downstream consequences. Instead, knockdown of VDAC1 or IP₃R in MERC-deficient cells abolished the hyperactive SOCE (**Figure 4F,G**). These experiments established the IP₃R-VDAC1 axis as the core machinery through which MERCs suppress SOCE. In this context, the SOCE-suppressing function of MERCs relies on IP₃R-VDAC1-mediated local Ca²⁺ transfer, rather than downstream consequences of mPTP for mitochondrial functions.

Finally, our studies unveiled a significant contribution of MT-EB1 comets to the tripartite STIM1 competition model regulating Ca^2+^ homeostasis. Previous studies have demonstrated that EB1, a microtubule plus-end tracking protein, binds and traps STIM1 at growing MT tips, thereby restricting its translocation to the plasma membrane (Chang *et al*., 2018). This prompted us to investigate whether MT-EB1 comets serve as the sink for surplus STIM1 released from disrupted MERCs. Indeed, upon MERC disruption, STIM1 was markedly enriched at EB1-labeled MT plus-ends (**Figure 6B**), indicating that MT-EB1 comets actively sequester the excess STIM1. Crucially, two orthogonal approaches that disrupt MT-EB1-mediated STIM1 sequestration without broadly altering ER morphology—EB1 knockdown and acute Spastin-mediated MT severing—both unmasked SOCE overactivation specifically in MERC-deficient cells (**Figure 6H** and **6I**), confirming that the MT-EB1 comets suppress SOCE through STIM1 sequestration.

The key feature of this MT-EB1 buffer is that it possesses a finite sequestration capacity. Two stoichiometric experiments further define the operational limits of this buffering system. First, overexpressing STIM1 in MERC-deficient cells overwhelmed the MT-EB1 buffer and recapitulated hyperactive SOCE (**Figure 6J**)—demonstrating that when the total STIM1 pool exceeds the sequestration capacity of MT-EB1 comets, the surplus STIM1 in MERC-deficient cells inevitably reaches the plasma membrane.

Conversely, EB1 overexpression effectively suppressed SOCE in control cells, yet failed to rescue the elevated SOCE in MERC-deficient cells (**Figure S6J**)—revealing that when the primary MERC trap is absent, the liberated STIM1 pool is too large for even an augmented MT-EB1 sink to contain. Together, these findings reveal an exquisite balance between STIM1 availability and the combined sequestration capacity of MERCs and MT-EB1 comets, wherein disruption of either component is tolerated but their simultaneous failure leads to pathological SOCE overactivation.

Beyond their direct role in STIM1 sequestration, MTs contribute to the structural organization of the ER network (Zheng *et al*, 2022). STIM1 has been shown to directly bind EB1 and form comet-like accumulations at polymerizing MT ends, a process that promotes ER tubule extension through the tip attachment complex mechanism (Grigoriev *et al*., 2008). Consistent with this intimate relationship between STIM1, MTs, and ER architecture, we observed that MERC disruption led to an expansion of ER-PM junctions (**Figure S3H**), suggesting that the loss of mitochondrial tethering redistributes ER membrane toward the cell cortex. The modest reduction of SOCE observed in control cells upon colchicine treatment (**Figure 6F**) may partly reflect secondary structural consequences of prolonged MT depolymerization on peripheral ER architecture, as extended absence of MTs has been shown to cause retraction of peripheral ER tubules away from the cell cortex (Terasaki *et al*, 1986; Waterman-Storer & Salmon, 1998). However, ER structural changes are unlikely to be the primary explanation for the phenotypes we report: EB1 knockdown, which disrupts STIM1-MT interactions without altering ER morphology, and acute Spastin-mediated MT severing, which avoids the prolonged drug exposure associated with colchicine, both consistently unmask SOCE upregulation or enhance STIM1 puncta in MERC-deficient cells (**Figure 6H, I**). These orthogonal approaches collectively support a model in which STIM1 sequestration by the MT-EB1 comets, rather than structural ER remodeling, is the principal mechanism underlying MT-dependent SOCE regulation. More broadly, while our study focused on the MT-STIM1 interface, actin filaments have also been shown to regulate ER-mitochondria contact structure through INF2-mediated polymerization (Chakrabarti *et al*, 2017), suggesting that multiple cytoskeletal elements cooperate to maintain the organellar architecture required for Ca^2+^ homeostasis.

In summary, our findings establish the PM, mitochondria, and the microtubule cytoskeleton as a tripartite, hierarchically organized regulatory network that governs SOCE through spatial competition for STIM1. Spontaneous mitochondrial Ca^2+^ flickering, driven by IP_3_R-VDAC1-mediated flux at MERCs, functions not merely as a local signal but as a homeostatic rheostat, which continuously calibrates the pool of STIM1 available for plasma membrane engagement. The identification of STIM1 competition among the PM, mitochondria and microtubules as a previously underappreciated dimension of Ca²⁺ signaling regulation opens new avenues for understanding how cells maintain homeostatic resilience. Given that dysregulation of both SOCE and mitochondrial Ca²⁺ is implicated in neurodegeneration, cardiac dysfunction and tumor progression, the tripartite network represents a conserved mechanism whose failure precipitates disease-associated Ca²⁺ overload.

## SUPPLEMENTARY INFORMATION

**Video S1.** Spontaneous mitochondrial Ca^2+^ flickers, related to Figure 1A.

**Video S2.** Mitochondrial [Ca^2+^] along with mitochondrial-targeted GFP, related to Figure 1C.

**Video S3.** Rapamycin induced mitochondria-ER contacts, related to Figure S2E, S2F.

**Figure S1.**
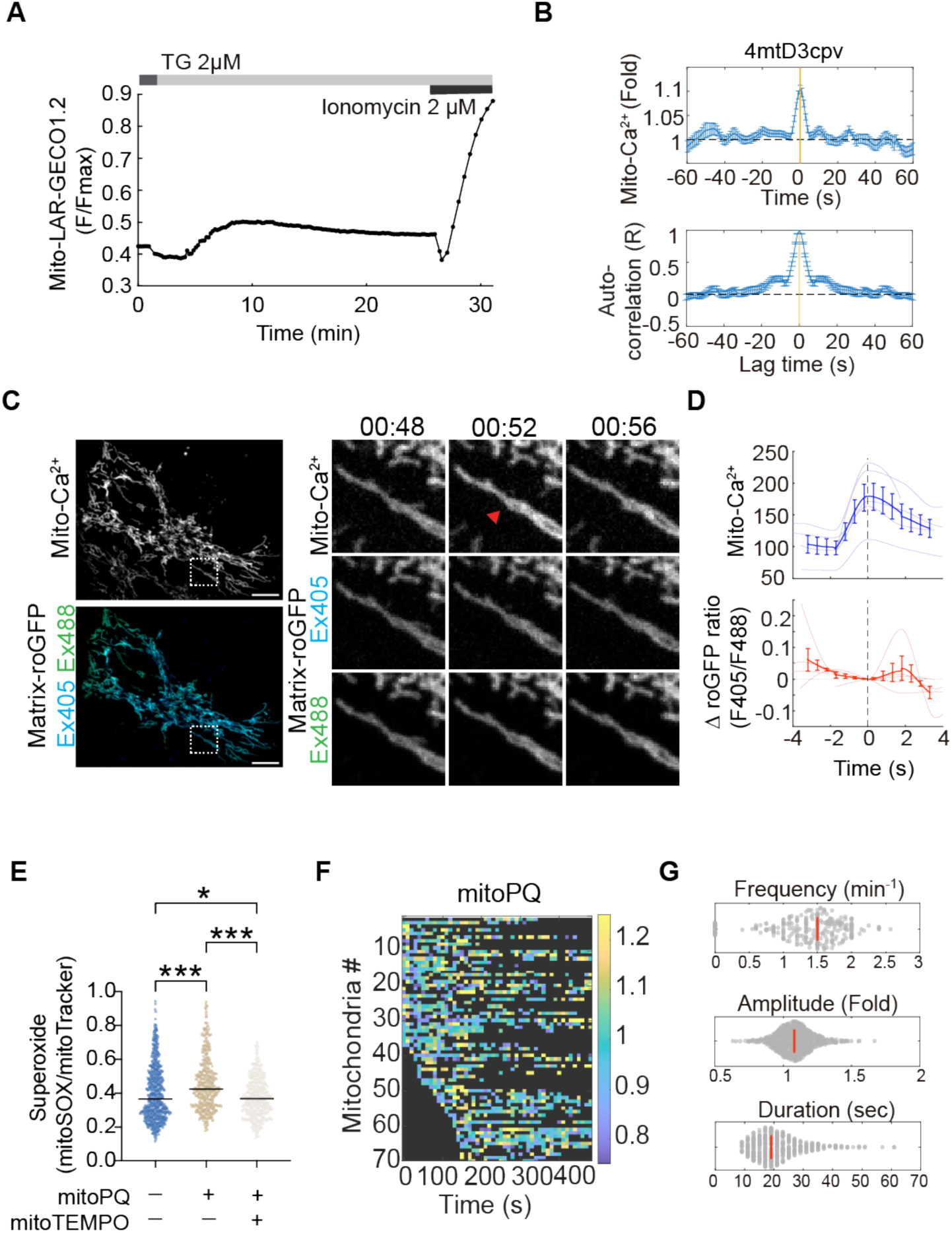
Validation of mitochondrial Ca^2+^ sensors and independence of Ca^2+^ flickers from redox status. **A.** Experimental protocol and validation of mito-LAR-GECO1.2 responsiveness. Cells were treated with thapsigargin (TG, 2 μM) followed by ionomycin (2 μM) to induce maximal mitochondrial Ca^2+^ influx. **B.** Peak-aligned average trace (top) and auto-correlation analysis (bottom) of the spontaneous Ca^2+^ transient signals measured by the FRET-based sensor 4mtD3cpv. **C.** Simultaneous live-cell imaging of mitochondrial Ca^2+^ and redox state. Representative confocal images of cells co-expressing mito-LAR-GECO1.2 (top, greyscale) and the dual-excitation ratiometric redox sensor matrix-roGFP (bottom, Ex405/Ex488). The red arrow indicates a spontaneous Ca^2+^ flicker event. Scale bars, 10 μm. **D.** Temporal alignment of the spontaneous mitochondrial Ca^2+^ flickers (top) with matrix-roGFP signals (bottom). For each detected Ca²⁺ flicker event, the roGFP ratio at each time point was normalized by subtracting the ratio value at the flicker peak (t = 0), yielding ΔF_405_/F_488_. The stable roGFP ratio trace confirms that Ca^2+^ events occur independently of transient redox fluctuations. **E.** Quantification of mitochondrial superoxide levels using MitoSOX (normalized to mitoTracker). Cells were untreated (control), treated with superoxide generator mitoPQ, or treated with mitoPQ plus ROS scavenger mitoTEMPO. Data are presented as a beeswarm plot, with horizontal lines indicating the median. Statistical analysis was performed using One-way ANOVA with Tukey’s post-hoc test. *p < 0.05, **p < 0.01, ***p < 0.001 **F.** Heatmap illustrating spatiotemporal dynamics of mitochondrial Ca^2+^ flickers measured by mito-LAR-GECO1.2 in cells treated with mitoPQ. **G.** Beeswarm plots showing the distribution of mitochondrial Ca^2+^ flicker frequency, amplitude, and duration under the mitoPQ treatment. Red lines indicate the median.

**Figure S2.**
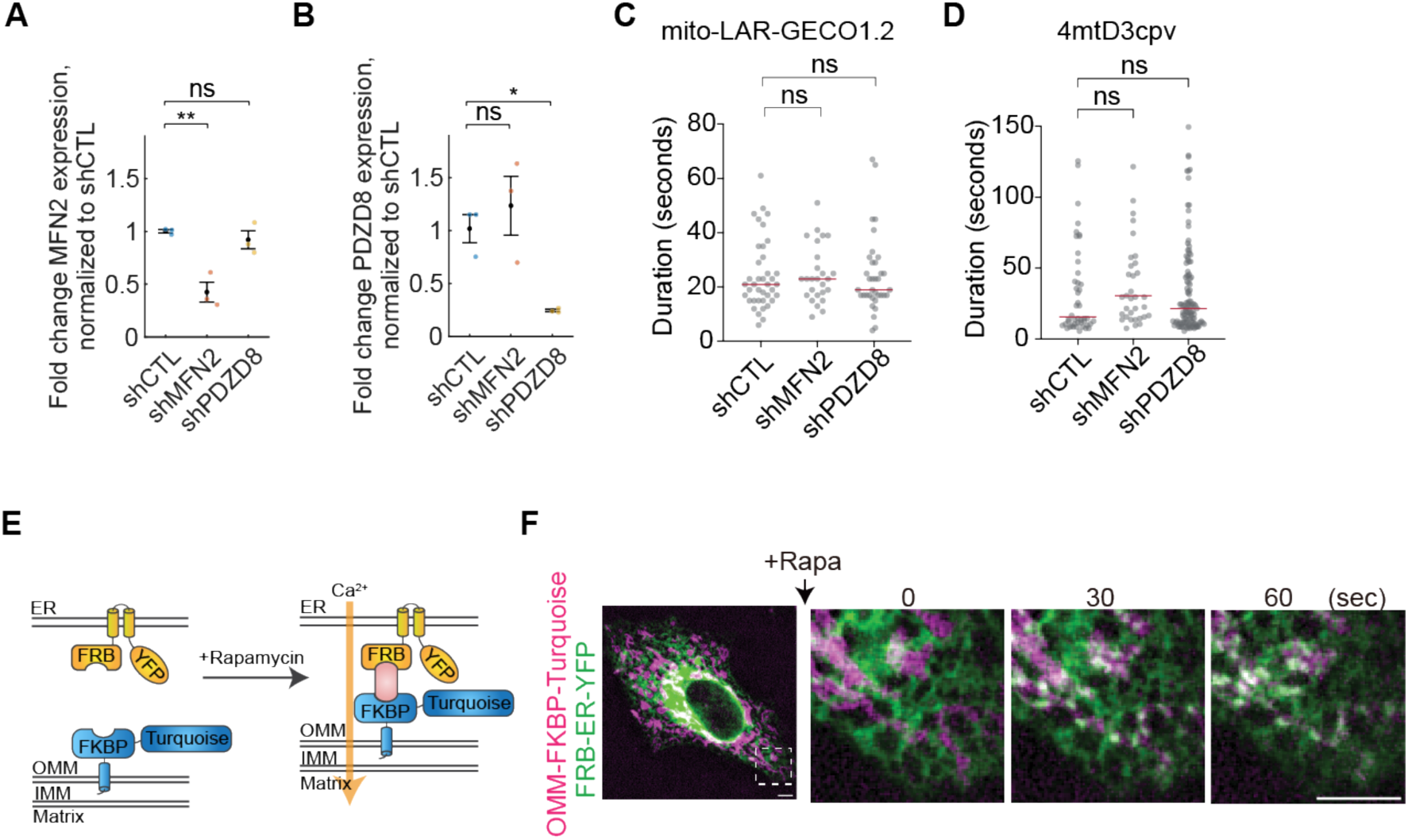
Validation of shRNA-mediated MERC knockdown, and FKBP–FRB-mediated MERC induction. **A–B.** RT-qPCR quantification of MFN2 (A) and PDZD8 (B) expression levels in control (shCTL) and respective knockdown (shMFN2 and shPDZD8) cells. Data are expressed as fold change relative to shCTL, normalized to HPRT. Data are presented as mean ± SEM. **C.** Quantitative analysis of mitochondrial Ca^2+^ flicker duration detected by mito-LAR-GECO1.2 in shCTL, shMFN2, and shPDZD8 cells. Red lines indicate the median. **D.** Quantitative analysis of mitochondrial Ca2+ flicker duration detected by 4mtD3cpv in shCTL, shMFN2, and shPDZD8 cells. Red lines indicate the median. **E.** Schematic of the rapamycin inducible synthetic tethering system. **F.** Representative time-lapse images of cells co-expressing OMM-FKBP-Turquoise (magenta) and FRB-ER-YFP (green) before and after rapamycin treatment (+Rapa). Enlarged insets highlight the rapid and progressive recruitment of the ER to mitochondria within 60 seconds. Scale bar, 5 µm. Statistical significance was determined by an unpaired, two-tailed Student’s t-test assuming equal variance for all panels (**p < 0.01).

**Figure S3.**
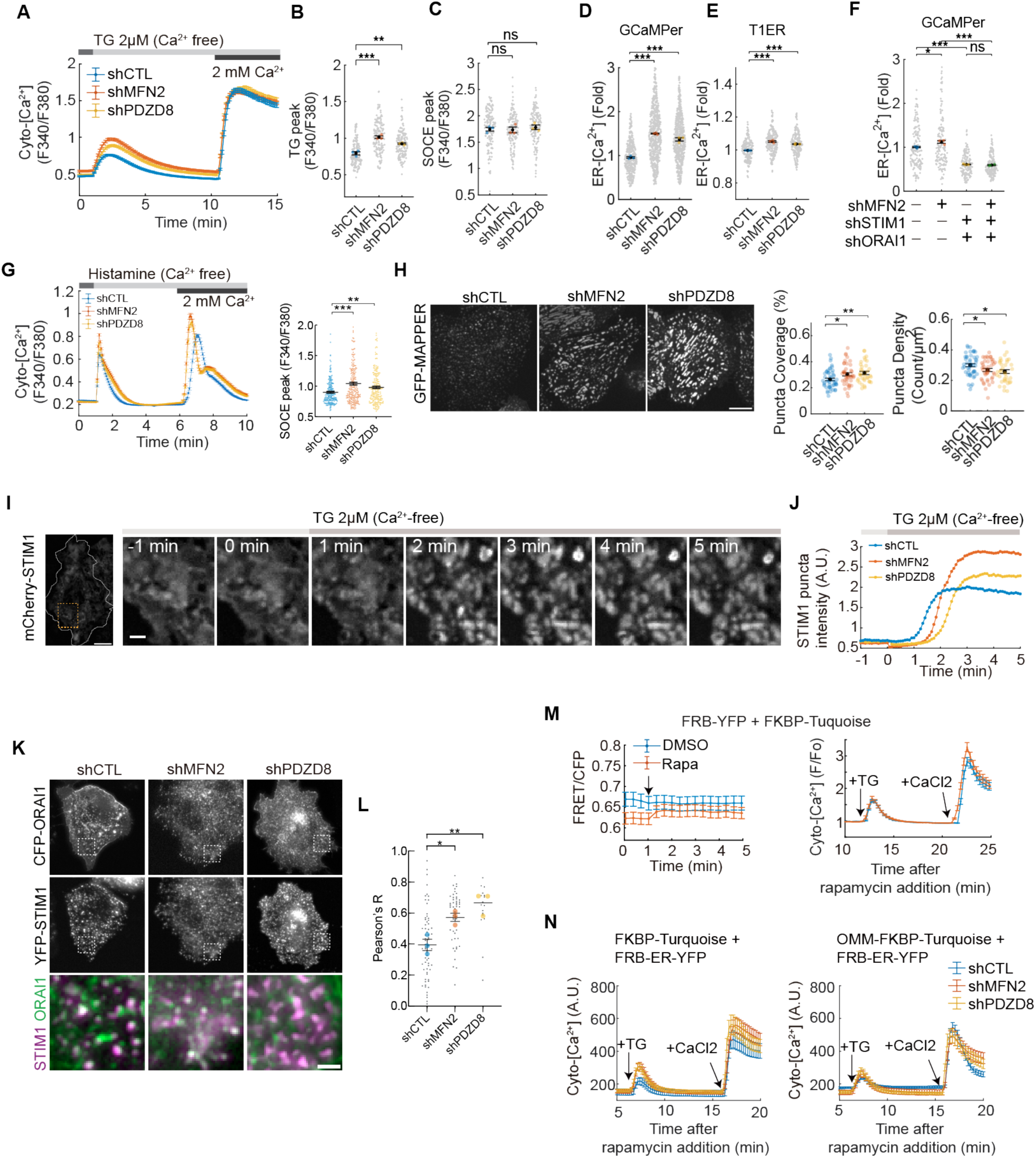
Functional and structural remodeling of the SOCE apparatus following MERC disruption. **A–C.** MERC disruption enhances ER Ca^2+^ release but not SOCE when microtubules are intact. (**A**) Representative Fura-2 traces of cytosolic Ca^2+^ changes in shCTL, shMFN2, and shPDZD8 cells in the absence of colchicine treatment. (**B, C**) Swarm charts quantifying the TG-induced Ca^2+^ release peak (**B**) and the subsequent SOCE peak (**C**). **D–E.** MERC disruption elevates basal ER Ca^2+^ levels. Quantification of the resting ER Ca^2+^ levels measured via ER-targeted Ca^2+^ sensors, GCaMPer (**D**) and T1ER (**E**). **F.** Hyper-filling of ER Ca^2+^ stores in MERC-deficient cells is STIM1–ORAI1-dependent. Swarm chart demonstrating that the elevation of resting ER Ca^2+^ level following MFN2 knockdown (shMFN2) is eliminated upon simultaneous knockdown of STIM1 (shSTIM1) and ORAI1 (shORAI1). **G.** MERC disruption enhances SOCE following physiological receptor-mediated stimulation. Representative Fura-2 traces (left) and quantification of SOCE peak amplitudes (right) in colchicine-pretreated shCTL, shMFN2, and shPDZD8 cells stimulated with histamine (100 µM), confirming that MERC-mediated SOCE suppression is not restricted to TG-induced store depletion. **H.** MERC disruption induces structural expansion of ER–PM junctions. Representative near-PM (basolateral) images of the ER–PM junction marker MAPPER in shCTL, shMFN2, and shPDZD8 cells (left panels), with corresponding quantification of MAPPER puncta coverage percentage (middle) and puncta density (right). Scale bar, 10µm **I–J.** Temporal dynamics of STIM1 puncta formation. (**I**) Representative time-lapse images demonstrating the rapid recruitment and clustering of mCherry–STIM1 following TG-induced store depletion. (**J**) Continuous traces showing the kinetics of STIM1 puncta intensity over time. Scale bar: 10 μm; inset, 1μm. **K–L.** MERC disruption promotes STIM1–ORAI1 colocalization at the plasma membrane. (**K**) Representative confocal images of cells co-expressing mCherry-STIM1 (magenta) and ORAI1-YFP. (green). (**L**) Quantification of STIM1–ORAI1 interaction using Pearson’s correlation coefficient (Pearson’s R). Scale bar: 1 μm **M.** Induction of cytosolic FKBP-FRB contacts does not suppress SOCE. (Left) FRET/CFP ratio traces confirming modest contact formation upon rapamycin addition in cells co-expressing cytosolic FRB-YFP and FKBP-Turquoise. (Right) Corresponding cytosolic Ca^2+^ traces confirming that neither the addition of rapamycin nor the formation of cytosolic FKBP-FRB contact suppresses SOCE, serving as a negative control for Figure 3K. **N.** Representative cytosolic Ca^2+^ traces corresponding to the quantitative swarm charts in Figure 3L. The traces illustrate SOCE responses following the rapamycin-induced formation of either non-specific contacts (left) or specific MERCs (right) across shCTL, shMFN2, and shPDZD8 cells. Quantitative data are presented as swarm charts with mean ± SEM. Small gray dots represent measurements from individual cells, while larger colored dots indicate the means of at least three independent replicates. Statistical analysis was performed using one-way ANOVA followed by Dunnett’s multiple comparisons test (*p < 0.05, **p < 0.01, ***p < 0.001).

**Figure S4.**
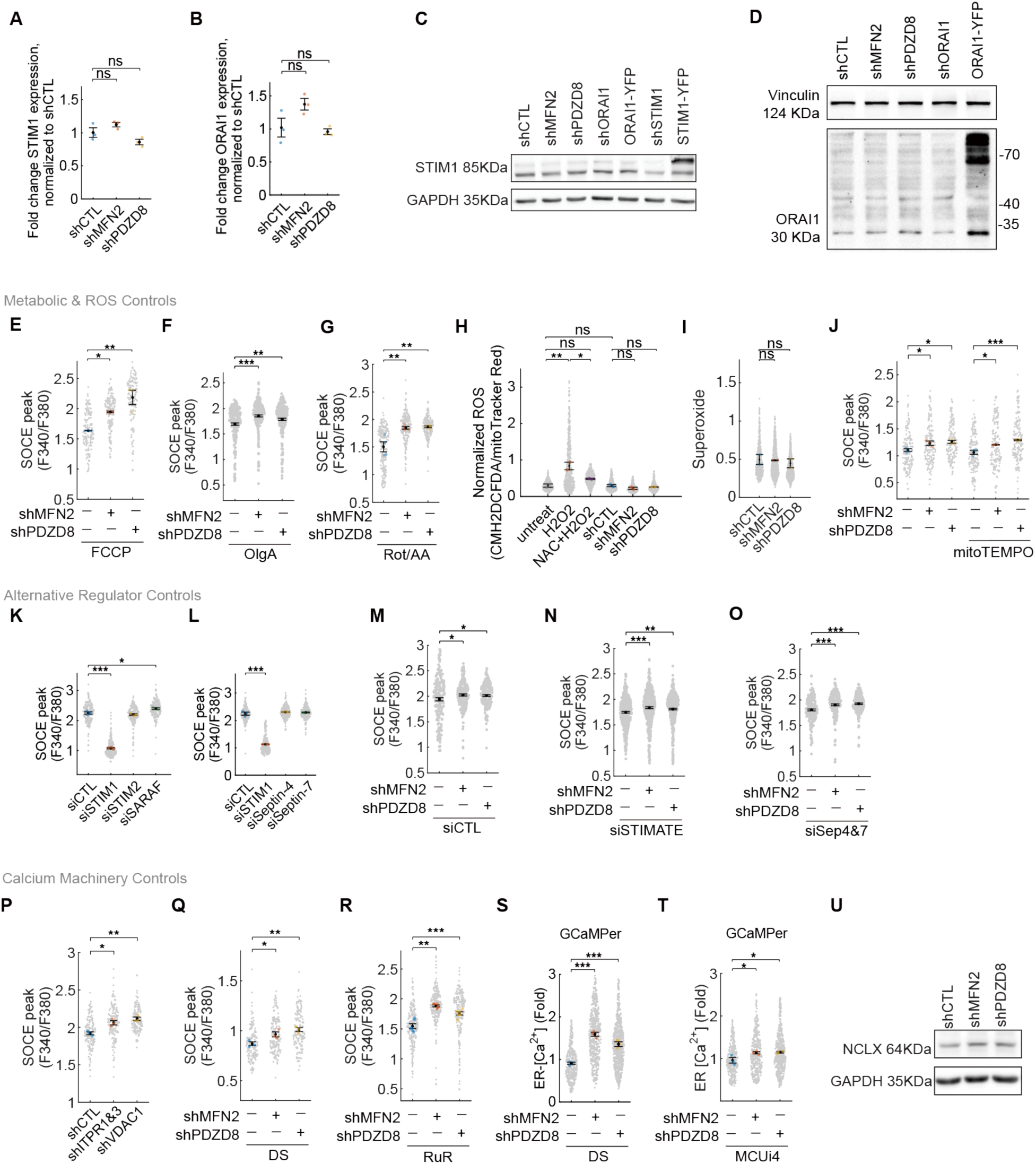
Mitochondrial metabolic outputs, ROS, and canonical STIM1 regulators do not account for MERC-mediated SOCE suppression. **A–D.** Expression levels of core SOCE components. mRNA levels of STIM1 (**A**) and ORAI1 (**B**), measured by RT-qPCR do not reveal significant alterations in shMFN2 and shPDZD8 compared to shCTL cells. Data are expressed as fold change relative to shCTL, normalized to HPRT. Western blots showing that protein levels of STIM1 (**C**) and ORAI1 (**D**) do not alter in shMFN2 and shPDZD8 compared to shCTL cells. These data confirm that MERC disruption does not alter the baseline expression of core SOCE machinery. **E–G.** Influence of mitochondrial metabolic perturbations. Quantification of SOCE peak amplitudes (F340/F380 ratio) in shMFN2 or shPDZD8 cells treated with the uncoupler FCCP (**E**), the ATP synthase inhibitor Oligomycin A (**F**), or the electron transport chain inhibitors Rotenone and Antimycin A (**G**). These metabolic perturbations fail to abolish SOCE upregulation induced by MERC disruption. **H–J.** Role of reactive oxygen species (ROS). (**H**) General cellular ROS levels measured by CM-H_2_DCFDA fluorescence normalized to MitoTracker Red. (**I**) Mitochondrial superoxide levels measured by MitoSOX. (**J**) Quantification of SOCE peaks showing that the mitochondrial ROS scavenger MitoTEMPO does not abolish the upregulated SOCE in shMFN2 or shPDZD8 cells. **K–O.** Influence of STIM1 regulators. Quantification of SOCE peak amplitudes in wild-type cells following siRNA-mediated knockdown of SARAF (**K**) or Septin-4 and Septin-7 (**L**). Notice that, compared to siCTL cells (M), knockdown of STIMATE (siSTIMATE, **N**) or Septin-4 and Septin-7 (siSep4&7, **O**) does not abolish the upregulated SOCE in shMFN2 or shPDZD8 cells. **P.** Phenotypic validation of the inter-organelle Ca^2+^ transfer machinery. Knocking down the ER Ca^2+^ release channels (IP_3_R1&3) or the outer mitochondrial membrane channel VDAC1 resulted in upregulated SOCE, phenocopying the effects of MERC disruption. **Q–R.** Effect of mitochondrial Ca^2+^ uptake inhibition on SOCE. MCU inhibitors DS16570511 (**Q**) or Ruthenium Red (**R**) fails to abolish the shMERC-induced SOCE upregulation. **S–T.** Measurement of ER Ca^2+^ content using ER-targeted sensor GCaMPer. Treatment with MCU inhibitors DS16570511 (**S**) or MCUi4 (**T**) fails to abolish the upregulated ER Ca^2+^ content in MERC-deficient cells. **U.** Western blot analysis of the mitochondrial Na^+^/Ca^2+^ exchanger (NCLX) in shMFN2 and shPDZD8 cells. Small gray dots represent measurements from individual cells, while larger colored dots indicate the means of at least three independent biological replicates. Error bars represent mean ± SEM. Statistical analysis was performed using one-way ANOVA followed by Dunnett’s multiple comparisons test (*p < 0.05, **p < 0.01, ***p < 0.001).

**Figure S5.**
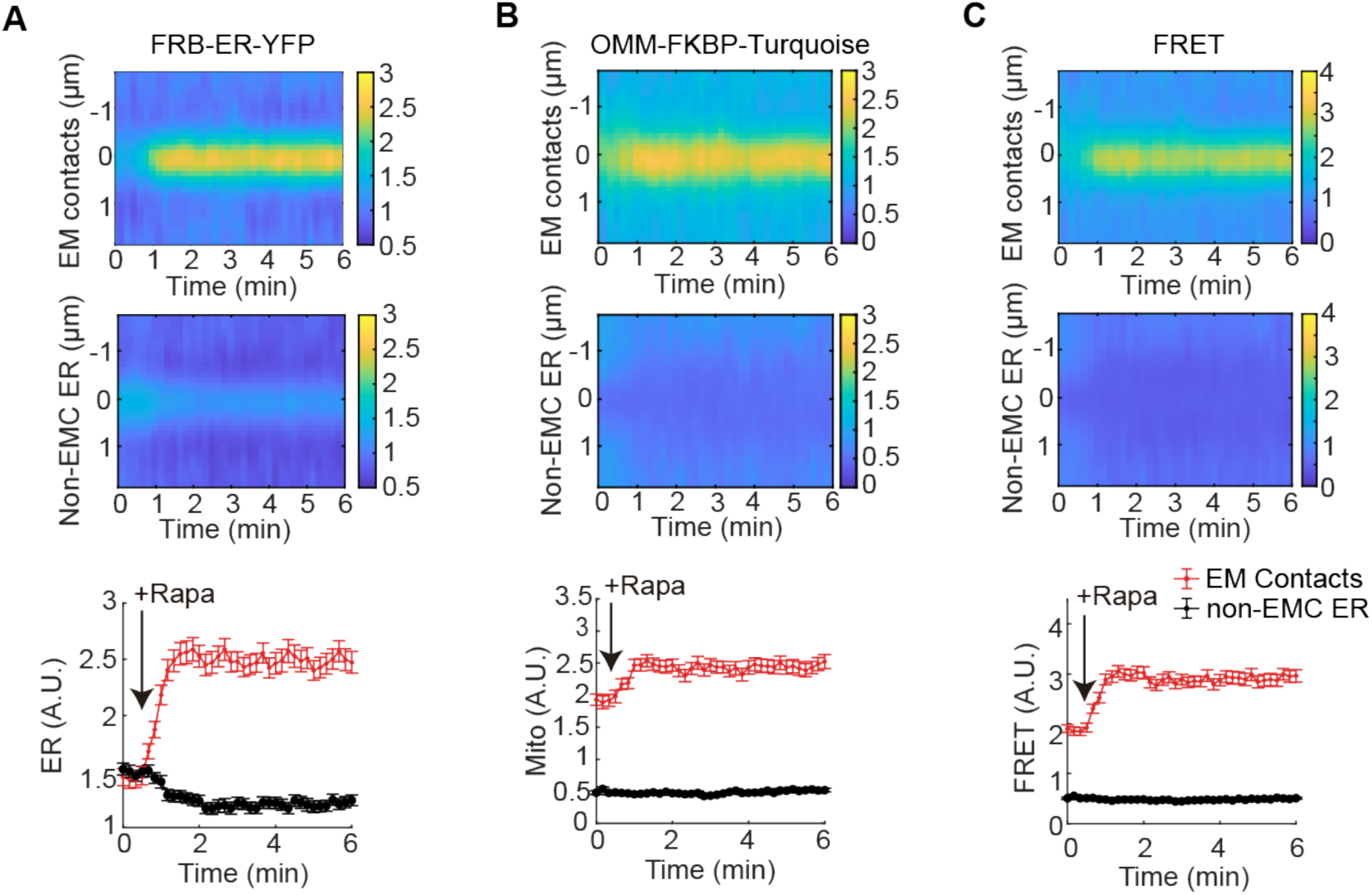
Kymograph analysis validates rapid and spatially restricted formation of induced MERCs. **A.** Spatiotemporal analysis of the ER surface marker. Kymographs illustrate the distribution of FRB-ER-YFP (ER surface marker) at induced MERCs (top) and non-contact ER regions (middle) over time. The bottom panel quantifies the rapid increase in FRB-ER-YFP intensity at contact sites (red) versus the reduction at non-contact regions (black) following rapamycin addition (arrow). **B–C.** Spatiotemporal analysis of the mitochondrial marker and tethering signal. Kymograph and line plot analyses of OMM-FKBP-Turquoise (B) and FRET signal (C) confirm the rapid and specific enrichment of mitochondrial tethers at contact sites following rapamycin addition. The lack of enrichment in non-contact regions (black traces and middle kymograph panels) demonstrates that the synthetic tethering system specifically modifies ER–mitochondria architecture without inducing global organelle redistribution. Data are represented as mean ± SEM.

**Figure S6.**
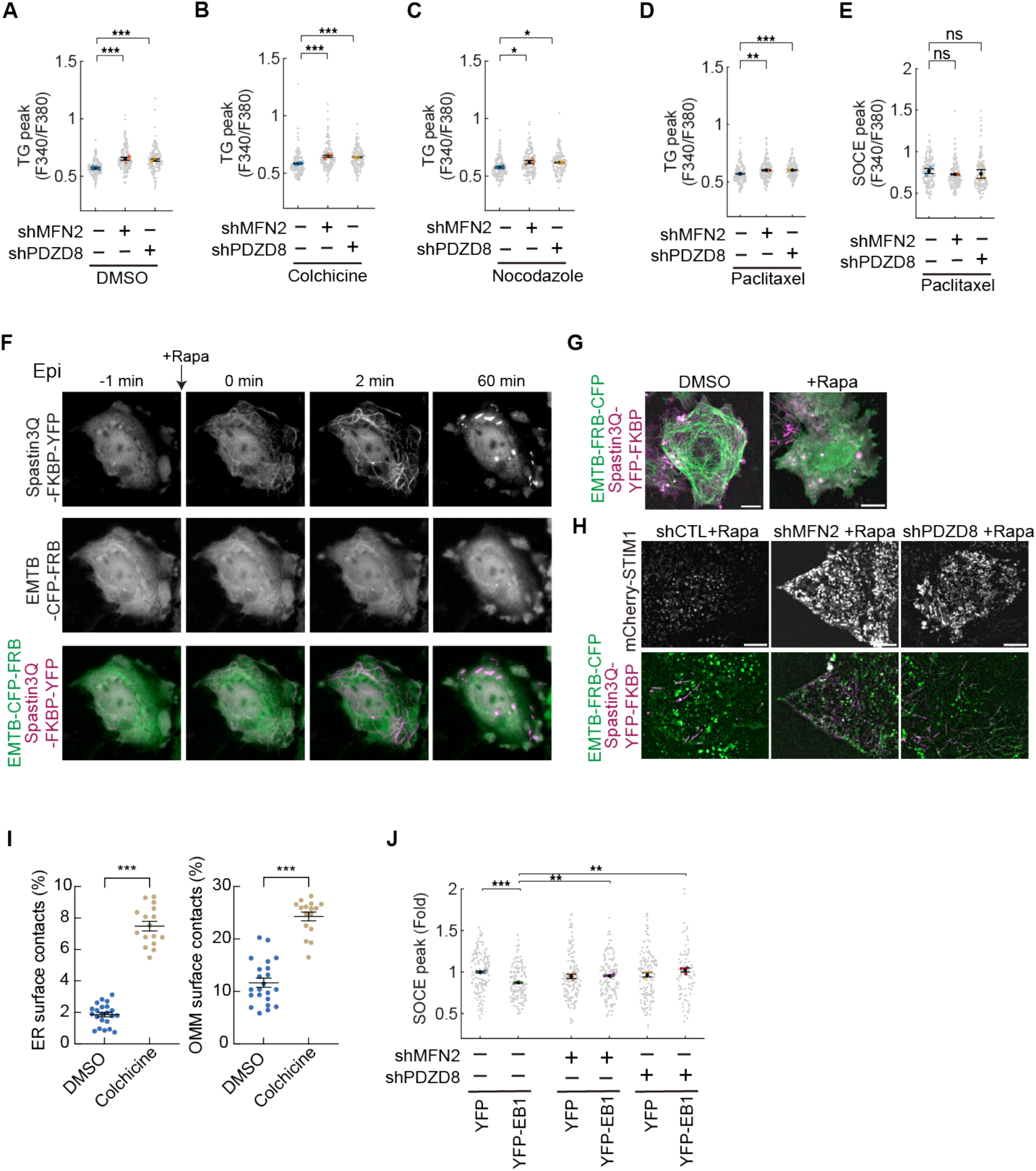
Validation of microtubule-mediated STIM1 regulation in MERC-deficient cells. **A–C.** Quantification of Thapsigargin (TG)-induced ER Ca²⁺ release in cells treated with vehicle (DMSO) (**A**), colchicine (**B**), or nocodazole (**C**). Microtubule disruption does not affect ER Ca²⁺ content in either control or MERC-deficient (shMFN2 or shPDZD8) cells. While microtubules contribute to SOCE regulation by sequestering STIM1 (see Fig. 6E**–G**), these results indicate that the pool of STIM1 required for maintaining basal ER Ca^2+^ homeostasis is not limited by microtubule-mediated sequestration. Given that the STIM1 requirement for basal ER Ca^2+^ maintenance is considerably lower than the demand during a maximal SOCE peak, these data suggest that microtubules have a finite capacity for STIM1 sequestration, allowing them to modulate SOCE without compromising basal ER Ca^2+^ storage. **D–E.** Measurements of (**D**) TG-induced ER Ca²⁺ release and (**E**) SOCE peak in cells treated with the microtubule stabilizer paclitaxel show that microtubule stabilization does not alter SOCE in MERC-deficient cells. **F.** Representative time-lapse images showing rapamycin-induced microtubule fragmentation using Spastin3Q–YFP–FKBP and EMTB–CFP–FRB. Spastin3Q–YFP–FKBP (top) is recruited to microtubules via the microtubule-binding protein EMTB–CFP–FRB (middle) upon addition of rapamycin (+Rapa). Bottom: merged images. Time 0 indicates the timepoint of rapamycin addition. Scale bar, 10 µm. **G.** Representative images showing the morphology of the microtubule network (green) and Spastin3Q localization (magenta) in DMSO vs. rapamycin-treated cells. Scale bar, 10 µm. **H.** Representative images of mCherry–STIM1 puncta (top) and microtubule disruption system (EMTB-FRB-CFP and Spastin3Q-YFP-FKBP, bottom) in control and MERC-deficient cells after rapamycin treatment. Scale bar, 10 µm. **I.** Microtubule disruption expands the extent of mitochondria–ER contacts. Quantification of the percentage of the ER surface in contact with mitochondria (left) and the mitochondrial outer membrane (OMM) surface in contact with the ER (right) in cells treated with vehicle (DMSO) or the microtubule-depolymerizing agent colchicine (10 μM) for 3 hours. Extent of contact was determined by 3D volumetric reconstruction of confocal Z-stacks. **J**. SOCE measurements in control and MERC-deficient cells expressing YFP or YFP–EB1. While EB1 overexpression reduces SOCE in control cells, it fails to suppress SOCE in MERC-deficient cells where STIM1 is not sequestered at MERCs. This suggests a limited capacity for EB1 to attenuate SOCE via STIM1 sequestration in the absence of intact MERCs. Data are presented as swarm charts with mean ± SEM; each dot represents an individual cell. Small gray dots represent measurements from individual cells, while larger colored dots indicate the means of at least three independent biological replicates. For J, statistical significance was determined by an unpaired, two-tailed Student’s t-test. For all other panels, statistical analysis was performed using one-way ANOVA followed by Dunnett’s multiple comparisons test (*p < 0.05, **p < 0.01, ***p < 0.001).

## METHOD

### Cell lines

HeLa cells and HepG2 cells were used for all the experiments. HEK293T cells were used to package all the lentivirus. These cells were cultured in humidified 37°C with 5% CO2 on culture dishes. The culture medium consisted of DMEM supplemented with 10% FBS and 1% penicillin/streptomycin. Cells were passaged upon reaching 90% confluence and used for up to 10 passages before a new aliquot was thawed.

### Pharmacological treatments

For scavenging ROS and mitochondria superoxide, cells were pretreated with 5 mM N-acetylcysteine (NAC) for 2 hours and 20 µM mitoTEMPO for 1 hour before assays. To induce mitochondria-specific superoxide, cells were incubated with 2 µM mitoPQ for 2 hours before assays. To disrupt microtubule polymerization, cells were pretreated with 10 μM colchicine or 1 μM nocodazole for 3 hours at 37°C prior to assays. DMSO was used as a vehicle control. For blocking mitochondrial Ca^2+^ uniporter (MCU), Ruthenium red (RuR) 5μM, DS16570511 30μM and MCUi4 5μM were added to cells 20 minutes before fura-2 assay. To inhibit mPTP, cyclosporin A (CysA) at 10 μM was applied to cells 20 minutes prior to the fura-2 assay. For blocking mitochondrial functions, we pretreated cells with 5 μM oligomycin A (OlgA), 0.5 μM FCCP, 5 µM CCCP, 1 μM Rotenone/Antimycin A for 20 minutes, respectively. For FKBP-FRB oligomerization induction, rapamycin 100 nM was added to the cells.

### DNA constructs

Rapamycin-induced oligomerization assay: To target FRB to the surface of ER, the C-terminus 521-587 sequence of *SAC1ML* was used. We first inserted FRB segment into YFP.N1 vector and fused the ER-target sequence between FRB segment and YFP segment using infusion cloning kit. To target FKBP to the outer membrane of mitochondria, the N-terminal 33 amino acids of *TOMM20* was used. We first fuse FKBP to Turquoise.N1 to generate a FKBP-Turquoise plasmid. After amplifying the OMM target sequence from human cDNA, the target sequence was further inserted to the N terminal FKBP-Turquoise.

MFN2 and PDZD8 constructs: *MFN2* and *PDZD8* were both amplified from human cDNA of HepG2 cells. Either MFN2 or PDZD8 sequence was inserted to N-terminal of p2A fragment followed by EYFP sequence. The MFN2-p2A-YFP and PDZD8-p2A-YFP sequence was further cloned into a lentiviral vector (pLAS2W.bsd).

STIM1 and ORAI1 constructs: YFP-STIM1 and their mutants (ΔK, D76A) were previously described (wild-type STIM1 Addgene #18857, STIM1 ΔK: Addgene #18861; STIM1 D76A: Addgene #18859). To use them with rapamycin-tethering system, we replaced all YFP with mCherry. 4mtD3cpv (Addgene #36324) and GFP-MAPPER (Addgene #117721) were bought from Addgene. mCherry-MAPPER was derived from GFP-MAPPER by using infusion cloning to replace the GFP sequence.

Plasmids for lentiviral production: RGECO1 (Addgene #32444), T1ER (Addgene #47928), mito-LAR-GECO1.2 (Addgene #61245), ER-LAR-GECO1 (Addgene #61244), GCaMPer (Addgene #65227) and GCaMP6s (Addgene #40753) have been previously described. To generate plasmids for lentiviral transduction, the coding region sequence of these plasmids were further cloned into lentiviral vectors (pLAS2w.bsd or pLAS2w.hyg from Academia Sinica, Taiwan).

### Lentiviral packaging and transduction

To knock down genes in cells, we used lentivirus to transduce shRNA into cells. shRNA plasmids were bought from National RNAi core (NRC; Academia Sinica, Taipei, Taiwan). HEK293T cells were plated on a 6-cm dish overnight until they reached ∼70% confluence. Three plasmids pCMV-D8.91, pMDG and shRNA plasmid, or cloned lentivector were transfected into cells with TransIT-LT1 (Mirus bio). After 16 hours transfection, medium was replaced with fresh DMEM with 1% bovine serum albumin (BSA) to remove the transfection reagent and DNA. Viral supernatant was collected at 48 hours and 72 hours after transfection. The collected viral supernatant was filtered through 0.45 µm polyethersulfone (PES) filter and then concentrated using Lenti-X concentrator (CloneTech, Takara Bio).

For gene knockdown experiments, lentiviral particles containing target-specific shRNAs were mixed and added to cells at MOI = 2 in culture medium supplemented with 8 μg/mL polybrene. Following infection, cells were selected by puromycin 2 µg/mL for 24 hours before being seeded into a collagen pre-coated 96-well plate for subsequent analysis.

### Stable cell lines establishment

Stable HeLa and HepG2 cell lines expressing genetically-encoded calcium sensors (R-GECO1, GCaMPer, T1ER, ER-LAR-GECO1, and mito-LAR-GECO1.2) were generated via lentiviral transduction. Transduced cells were selected using either Blasticidin (10 µg/mL) or Hygromycin (200 µg/mL). To ensure uniform expression levels, cells were further sorted by flow cytometry based on fluorescence intensity.

### Transfection

For plasmid transfections, HepG2 and HeLa cells were transiently transfected using Lipofectamine 3000 or TransIT-X2, respectively, according to the manufacturer’s instructions. Cells were used for subsequent experiments 12-16 hours post-transfection. For RNAi, cells were transfected with negative control or gene-specific siRNA oligos following the manufacturer’s protocol. After 16 hours, the transfection medium was replaced with fresh medium, and cells were used for the following experiment at 48-72 hours post-transfection. To minimize potential Ca²⁺ buffering effects associated with stable sensor expression, transient transfection was used for the following experiments: (i) all 4mtD3cpv-based and MCU inhibition mitochondrial Ca²⁺ measurements (Figures 1G–1N and 2G–2I); (ii) ER Ca²⁺ measurements using GCaMPer (Figures S3D), T1ER (Figure S3E) and ER-LAR-GECO1 (Figure 5A).

### Ca^2+^ imaging

#### Mitochondrial [Ca^2+^] measurement

Mitochondrial [Ca^2+^] was detected using both mito-LAR-GECO1.2 and 4mtD3cpv. The stable cells expressed mito-LAR-GECO1.2 were seeded into an 8-well chamber slide and washed with ECB twice before imaging. For detecting mitochondrial Ca^2+^ with 4mtD3cpv, HeLa cells were seeded in 8 well chamber slide overnight followed by transient transfection.

Time-lapse images were acquired at 5-second intervals for 10 minutes using a spinning disc confocal microscope (Nikon Eclipse Ti equipped with a Yokogawa CSU-X1 spinning disc unit) with a 60x oil immersion objective (CFI Plan Apo Lambda, NA 1.4). Images were captured using 561 nm laser excitation and 590/50 nm emission filter.

The time-lapse images were processed for segmentation and tracking using the MATLAB algorithm mitometer (Lefebvre *et al*, 2021). For each mitochondrial track, the fluorescence intensity was normalized to its mean intensity across time (F/Fmean). Mitochondrial Ca^2+^ flickers events were identified using local maxima detection with prominence thresholding (islocalmax function), followed by interpolation with a 10-fold increase in temporal resolution for precise alignment and analysis. The frequency of Ca^2+^ flickers was quantified as the number of detected peaks per minute, while the amplitude was measured as the normalized fluorescence intensity at peak positions. For temporal pattern analysis, individual Ca^2+^ flicker events were aligned to their peak positions (t=0). Event periodicity was assessed using autocorrelation analysis, with flicker duration determined from the temporal width of autocorrelation peaks. The temporal distribution of events was further analyzed by comparing the observed frequency distribution to theoretical Poisson statistics. Statistical comparisons between experimental groups were performed using the student’s t-test method.

### SOCE activity assay

HeLa cells were plated into a 96-well plate 24-48 hours before imaging. For experiments involving depolymerized microtubules, cells underwent a pre-treatment with 10 µM colchicine for three hours to ensure effective microtubule disruption. Cells were loaded with Fura-2, AM at a concentration of 2 µM, supplemented with 0.1% Pluronic F127 (Invitrogen) and 1 µM probenecid (Invitrogen) in extracellular buffer (ECB composition: NaCl 125mM, KCl 5mM, MgCl_2_ 1.5mM, Glucose 10mM, CaCl_2_ 1.5mM and HEPES 20mM) at room temperature for 20 minutes. After loading, cells were washed with ECB twice before imaging. Imaging was then carried out using a 4x objective (Nikon CFI S Fluor 4x) on a Nikon Eclipse Ti microscope. Baseline levels were recorded for 1 minute every 5-10 seconds. Thapsigargin (TG) 2 µM and EGTA 3 mM were added to induce calcium release from the endoplasmic reticulum (ER), with imaging continued for nine minutes post-addition to monitor transient calcium levels. Subsequently, 2mM CaCl_2_ was introduced to the system to assess store-operated calcium entry (SOCE), and imaging was extended for an additional five minutes. Relative calcium levels were quantified from the captured images using customized MATLAB scripts designed to analyze changes in fluorescence intensity.

### ER-[Ca^2+^] measurement

ER-[Ca^2+^] level was examined by either GCaMPer, T1ER or ER-LAR-GECO1. The stable cells were seeded into a 96-well plate 24-48 hours before imaging. Time-lapse imaging was performed using a Nikon Eclipse Ti microscope with a 20x objective (Nikon Plan Apo lambda D 20x). Baseline images were taken for 30 seconds every 5 seconds. TG was added to cells to induce ER-Ca^2+^ leakage and images were taken for subsequent 10 minutes. Subsequent analysis was performed using customized MATLAB scripts.

### 3D reconstruction and quantification of mitochondria-ER contact sites

For 3D reconstruction, z-stacks images acquired in the ER-Tracker Green / MitoTracker Red channels were taken with a spinning disc confocal microscope (Nikon Eclipse Ti equipped with a Yokogawa CSU-X1 spinning disc unit) with a 60x oil immersion objective (CFI Plan Apo Lambda, NA 1.4). To quantify MERCs, we used a semi-automated 3D image-processing pipeline in MATLAB. Individual cells from z-stack images were isolated using a semi-automated ROI tool. To optimize spatial resolution, cropped 3D stacks underwent customized background subtraction function (rolling-ball method), Gaussian smoothing and Richardson-Lucy deconvolution using 3D Gaussian PSFs tailored to experimental parameters (NA = 1.4; axial step size = 0.3 µm ). Following deconvolution, 3D volumes were segmented via adaptive thresholding, size-exclusion filtering (minimum volume = 50 voxels), and morphological hole-filling. The 3D outer boundaries (surfaces) of both organelles were mathematically isolated by subtracting a morphologically eroded mask (1-voxel radius) from the original binary volume. 3D volumetric contact sites were computed via the spatial intersection of the fully segmented ER and mitochondrial volumes. The extent of physical interaction was calculated as the percentage of the total organelle surface area directly overlapping with the adjacent organelle’s boundary. 3D profiles were visually verified using isosurface patch reconstructions.

### Live-cell imaging Redox detection

HeLa cells co-transfected with matrix-roGFP and CMV-LAR-GECO1.2 were seeded into a 8-well chamber slide. The culture medium was replaced with ECB containing 40mM HEPES prior to imaging. Time-lapse roGFP images were captured using both 405 nm and 488 nm laser excitation and 520/35nm emission filter. The Mito-[Ca^2+^] signal was simultaneously recorded as described above.

### ROS detection

To assess general cellular ROS levels, cells were incubated with 5 µM CH-H2DCFDA (Invitrogen, C6827) and 100nM MitoTracker Red CMXRos (Invitrogen, M46752) in ECB for 20 minutes at 37°C in the dark. Following incubation, cells were washed with ECB twice and immediately imaged. CM-H_2_DCFDA fluorescence was acquired using 488 nm laser excitation with a 520/35 nm emission filter, and MitoTracker Red CMXRos was acquired using 561 nm laser with 590/50nm emission filter. ROS levels were quantified as the ratio of CM-H_2_DCFDA fluorescence intensity normalized to MitoTracker Red intensity to correct mitochondrial mass.

To assess mitochondrial superoxide levels, cells were incubated with 5 μM MitoSOX Red (Invitrogen, M36008) and 100 nM MitoTracker Green FM (Invitrogen, M7514) in ECB for 20 minutes at 37°C in the dark, protected from light to prevent photooxidation. Mitochondrial superoxide levels were quantified as MitoSOX Red fluorescence intensity normalized to MitoTracker Green intensity to correct for mitochondrial mass.

### STIM1 dynamics

Stable HeLa cells expressing mCherry-STIM1 were knocked down using shRNA lentivirus and seeded into an 8-well chamber slide. Cells were pre-treated with either DMSO or 10 µM colchicine in culture medium and incubated at 37°C. After 3 hours, the culture medium was replaced with ECB containing 40 mM HEPES and either DMSO or 10 µM colchicine. Time-lapse imaging was performed using a spinning disc confocal microscope (Nikon Eclipse Ti equipped with a Yokogawa CSU-X1 spinning disc unit) and a 60x oil immersion objective (CFI Plan Apo Lambda) at room temperature. The focal plane was set at the bottom of the cells, and images were captured every 5 seconds. One minute after the start of imaging, 2 µM thapsigargin (TG) and EGTA were added to induce calcium (Ca²⁺) leakage from the endoplasmic reticulum (ER). Imaging continued immediately after the addition of these agents, with the dynamic responses of STIM1 monitored for an additional 10 minutes.

### Mitochondria-ER contacts inducing system

HeLa cells, or stable HeLa cells expressing the cytosolic Ca²⁺ indicator R-GECO1, were used for the expression of FRB-ER-YFP and Mito-FKBP-Turquoise. The cells were transfected with plasmids encoding FRB-ER-YFP and Mito-FKBP-Turquoise 24 hours before the assay. Before imaging, the cells were washed twice, and the medium was replaced with ECB. To induce oligomerization between FRB and FKBP, 100 nM rapamycin was added at the indicated time in the experiment. Mitochondria-ER contacts were monitored using FRET.

### Microtubule erasers

Stable HeLa cells expressing mCherry-STIM1 were used for the expression of EMTB-CFP-FRB and Spastin3Q-YFP-FKBP. 100nM rapamycin was added to cells to recruit dSpastin onto microtubules. Images were taken every 10 minutes for 1 hour to visualize the degradation of microtubules. For STIM1 puncta analysis, rapamycin was added to cells for 5 minutes to degrade microtubules, followed by treatment with TG for 10 minutes to induce STIM1 puncta formation for subsequent analysis.

### Immunofluorescence staining

For EB1 staining, HeLa cells were transfected with mCherry-STIM1 using the TransIT-X2 transfection reagent (Mirus Bio) for 12–16 hours prior to immunostaining. Following transfection, the cells were rinsed with PBS and fixed in methanol at −20°C for 10 minutes. The cells were then washed twice with PBS at room temperature, with each wash lasting 5 minutes. Subsequently, the cells were blocked in 1% BSA in PBS for 1 hour at room temperature. After blocking, cells were incubated overnight with the primary antibody at 4°C (Anti EB1, Abcam ab53358). The following day, the cells were washed twice with PBS for 5 minutes at room temperature. The secondary antibody, Alexa Fluor 488 (Invitrogen #A48262TR), was applied to stain EB1, and DAPI was used to counterstain nuclear DNA. Imaging was conducted using a spinning disc confocal microscope (Nikon Eclipse Ti equipped with a Yokogawa CSU-X1 spinning disc unit) and a 100x oil immersion objective (CFI Plan Apo Lambda).

### Western blot

HepG2 cells were utilized to assess the expression levels of STIM1 and ORAI1. The cells were treated with shRNAs targeting MFN2, PDZD8, STIM1, and ORAI1, as well as STIM1-YFP and ORAI1-YFP constructs. Following puromycin selection, cells were lysed in RIPA buffer (25 mM Tris-HCl, pH 7.6, 150 mM NaCl, 1% Triton X-100, 1% sodium deoxycholate, 0.1% SDS) supplemented with protease inhibitor cocktail and incubated on ice. The lysates were centrifuged at 15,000g for 15 minutes, and the supernatants were collected to quantify protein concentrations. For SDS-PAGE, 30 µg of each protein sample was mixed with an equal volume of sample buffer and boiled at 95°C for 5 minutes. The samples were then loaded onto a 10% SDS-PAGE gel for electrophoresis. After separation, the proteins were transferred onto a polyvinylidene difluoride (PVDF) membrane. The membrane was blocked in 3% BSA in Tris-buffered saline with 0.05% Tween-20 (TBST) for 1 hour at room temperature, followed by overnight incubation at 4°C with the following primary antibodies: anti-STIM1 (Cell Signaling Technology, #5668) at 1:1,000, anti-ORAI1 (Abcam, ab111960) at 1:500, anti-NCLX/SLC24A6 (Proteintech, #21430-1-AP) at 1:1,000, anti-GAPDH (Cell Signaling Technology, #2118S) at 1:10,000, and anti-Vinculin (Cell Signaling Technology, #13901S) at 1:10,000. The membranes were then incubated with appropriate secondary antibodies (Sigma-Aldrich AP182P or AP192P) at 1:10,000 for 1 hour at room temperature. Enhanced chemiluminescence (T-Pro Biotechnology, New Taipei City, Taiwan) was used to detect the proteins, and the chemiluminescent signals were visualized using the Bio-Rad Gel Doc2000 system (Bio-Rad, Hercules, CA, USA).

### RNA extraction and expression analysis

Total RNA was isolated from HepG2 cells using NucleoZol reagent (MACHEREY-NAGEL, Düren, Germany) following the manufacturer’s instructions. The isolated RNA was subsequently reverse-transcribed into complementary DNA (cDNA) using SuperScript IV Reverse Transcriptase (Invitrogen). Quantitative real-time PCR (qPCR) was performed to evaluate gene expression levels using SYBR Green Master Mix (Bio-Rad). The qPCR reactions were prepared in a 96-well plate (Bio-Rad) and analyzed on the CFX Connect Real-Time PCR Detection System (Bio-Rad). Gene expression was normalized to HPRT as the internal reference gene using the 2^−ΔΔCt^ method.

### Quantification and statistical analysis

All Ca^2+^ imaging and FRET data were analyzed using customized MATLAB scripts (MATLAB r2024a, Mathworks). For cytosolic Ca^2+^ measurements using Fura-2, the ratio of fluorescence intensities at 340 nm and 380 nm excitation (F340/F380) was calculated after background subtraction at each wavelength, presented as arbitrary units (A.U.). For mitochondrial Ca^2+^ flicker analysis, fluorescence intensities were normalized to the temporal mean values of individual mitochondria and presented as fold changes. For other Ca^2+^ measurements, background-subtracted fluorescence intensities were either presented directly as intensities (A.U.) or normalized to baseline (F/F0) depending on the experimental context. For FRET analysis, background-subtracted FRET signals were normalized to donor (CFP) intensities to calculate FRET/CFP ratios. When comparing between experimental groups, data were sometimes normalized to control conditions and presented as fold changes relative to control.

Quantitative data are presented as mean ± standard error of the mean (s.e.m.) unless otherwise specified in figure legends. For swarm plot quantifications, small gray dots represent measurements from individual cells, and larger colored dots indicate the means of at least three independent biological replicates; statistical comparisons were performed on the biological-replicate means. The specific statistical test used for each panel is indicated in the corresponding figure legend. In brief, two-group comparisons were performed using an unpaired, two-tailed Student’s t-test (assuming equal variance) or, where data did not meet the assumptions of parametric testing, the non-parametric Mann–Whitney U test. Multi-group comparisons were performed using one-way ANOVA followed by Dunnett’s multiple comparisons test (when each group was compared with a single control) or Tukey’s multiple comparisons test (when all pairwise comparisons were assessed).

Categorical data were analyzed using the Chi-square test. The relationship between STIM1 recruitment to EB1 comets and total cellular STIM1 expression was analyzed using zero-intercept ANCOVA, with the regression slope (β) compared across groups. The stochastic nature of mitochondrial Ca²⁺ flickers was assessed by fitting the observed frequency distribution to a Poisson distribution. Statistical analyses were performed in MATLAB r2024a (MathWorks) and GraphPad Prism 10.3.1 (GraphPad Software). Statistical significance is denoted as follows: *p < 0.05, **p < 0.01, ***p < 0.001.

